# Human Polyomavirus-Encoded Circular RNAs

**DOI:** 10.1101/2020.12.22.423831

**Authors:** Rong Yang, Eunice E. Lee, Jiwoong Kim, Joon H. Choi, Yating Chen, Clair Crewe, Philipp E. Scherer, Elysha Kolitz, Clay Cockerell, Taylor R. Smith, Leslie Rosen, Louisa Verlinden, Mariet C. Feltkamp, Christopher S. Sullivan, Richard C. Wang

## Abstract

Circular RNAs (circRNAs) are a conserved class of RNAs with diverse functions. A subset of circRNAs are translated into peptides. Here we describe circular RNAs encoded by human polyomaviruses (HPyVs), including circular forms of RNAs encoding variants of the previously described alternative large T antigen open reading frame (ALTO) gene. Circular ALTO RNAs (circALTOs) can be detected in virus positive Merkel cell carcinoma (VP-MCC) cell lines and tumor samples. CircALTOs are stable, predominantly located in the cytoplasm, and N^6^-methyladenosine (m^6^A) modified. MCPyV circALTOs produce ALTO protein in cultured cells. MCPyV ALTO promotes the transcription of co-transfected reporter genes. MCPyV circALTOs are enriched in exosomes derived from VP-MCC lines and circALTO-transfected 293T cells, and purified exosomes can mediate ALTO expression and transcriptional activation. The related trichodysplasia spinulosa polyomavirus (TSPyV) also expresses a circALTO that can be detected in infected tissues and produces ALTO protein in cultured cells. Thus, human polyomavirus circRNAs are expressed in human tumors and tissues, encode for proteins, and may contribute to the infectious and tumorigenic properties of these viruses.

## INTRODUCTION

About a dozen polyomaviruses infect humans (1). Merkel cell carcinoma (MCC) is a rare, deadly skin cancer linked to Merkel cell polyomavirus (MCV or MCPyV) infection in 80% of cases (2). In addition, trichodysplasia spinulosa polyomavirus (TSPyV) causes a primary infection that presents as eruptions of follicular papules and spicules in immunosuppressed patients (3-5). The small, double-stranded genome of polyomaviruses includes early, late, and non-coding control regions (6). The polyomavirus early region (ER) encode for small T-antigen (sT) and large T-antigen (LT), while the late region contains late structural proteins VP1 and VP2 (7-9). Moreover, MCPyV has been reported to encode a microRNA (miRNA) that lies antisense to the early transcripts (10,11). In a subset of *Alphapolyomaviruses*, including MCPyV and TSPyV, the early region has been shown to generate several variably spliced transcripts, including a transcript that encodes a protein in an alternate reading frame from large T antigen (ALTO) (12,13). ALTO shares a common evolutionary origin with middle T antigen (MT) of murine polyomavirus (MPyV) (12,14). Although MPyV MT has transforming activity and has been a useful tool to study cellular transformation (15), the precise function of human ALTO proteins remain unclear.

Circular RNAs (circRNAs) are covalently closed single-stranded RNAs that are most commonly produced by non-canonical 3’-5’ ligation or splicing of linear RNA (16,17). More than 40 years ago, viroids were the first recognized single-stranded circular RNA species (18). Later, RNAs with circular structure were observed in eukaryotic cell lines by electron microscopy (EM) (19), and were then found in all major domains of life (20,21). Recent improvements in high-throughput RNA sequencing and computational approaches have revealed circRNAs to be regulated and abundant (22,23). In addition to the endogenous circRNAs generated by eukaryotic cells, an increasing number of viruses have been found to encode for circRNAs. Epstein barr virus (EBV), Kaposi’s sarcoma herpesvirus (KSHV), and human papillomaviruses (HPVs) have all been demonstrated to generate circRNAs (24-28). The HPV-derived circRNA, circE7, encodes for the E7 oncoprotein and is essential for the transformed growth of CaSki cervical carcinoma cells (29).

CircRNAs exert their functions in eukaryotic cells through a variety of mechanisms. They were first recognized to act as miRNA sponges (30,31), such as ciRS-7/CDR1as, which acts as a miRNA sponge or decoy for miR-7 (31). Some circRNAs bind to RNA-binding proteins (e.g. HuR, RNA Pol II) to influence transcription and translation (32,33). Notably, numerous circRNAs can be translated to functional proteins (29,34). CircRNAs have been found to be resistant to degradation and also enriched in extracellular vesicles, including exosomes and virions, suggesting ways in which circRNAs could differ from their more abundant linear counterparts to exert unique biological functions (27,35,36).

Here we describe the identification of circular RNAs encoded by human polyomaviruses. MCPyV has two circRNAs in the early region encompassing the ALTO gene (circALTO1 and circALTO2) that could be detected in VP-MCC cell lines and patient tumors. TSPyV encodes a smaller circALTO, which could be efficiently generated in vitro and detected in infected tissues from TS patients. Both MCPyV and TSPyV circALTOs encoded for ALTO in vitro. Moreover, MCPyV circALTOs could be detected in the exosomes of VP-MCC cells, and circALTO-containing exosomes could mediate ALTO expression in recipient cells. Finally, MCPyV circALTOs-mediated ALTO expression increased the expression of co-transfected reporter genes. This study identifies and characterizes the first known polyomavirus encoded circular RNAs and demonstrates the first known function for HPyV ALTO, suggesting novel ways in which circRNAs could influence the infectious and transforming activities of HPyVs.

## MATERIALS AND METHODS

### De novo circular RNA detection from circular viral genomes

Identification of circular RNA from RNA sequencing (RNA-Seq) data of viruses with circular genomes was essentially described previously (29). In brief, a custom pipeline termed vircircRNA was used (https://github.com/jiwoongbio/vircircRNA) and Burrows-Wheeler Aligner (BWA, v0.7.15) were used for aligning RNA-Seq reads. Splice reads mapped in chiastic order were defined as back-splice reads from circular RNAs.

### Circular RNA detection from public RNA-Seq data

The genome sequences of MCPyV (NC_010277.2) were downloaded from National Center for Biotechnology Information (NCBI) database. We searched public sequencing data of MCPyV from NCBI Sequence Read Archive (SRA). Backsplice junctions were detected from the following SRA samples: ERS760222-5 (37).

### Cell Culture

MKL1, MKL2, MS-1 and WaGa cells were grown in RPMI supplemented with 20% fetal bovine serum (FBS). MCC13, MCC26, and UISO cells were cultured in RPMI supplemented with 10% FBS. MCC lines were confirmed through verifying the expression of sT and assessment of cell morphology. Characteristics of these cell lines have been described previously (7,38). WaGa shows high expression of MCPyV early region genes (7,39). HeLa, HEK293 and HEK293T cells (ATCC) were cultured in DMEM and additional 10% FBS. All cells were grown in incubator at 37°C with 5% CO2.

### End-point PCR, qRT-PCR and RNase R treatment

End point PCR and quantitative real-time RT-PCR (qRT-PCR) for circRNAs were performed following the protocol described previously (29). In brief, total RNA was extracted from cells using the RNeasy Mini Kit (Qiagen, 74104). 2 μg of total RNA was subjected to RNase R treatment (29) with 5U RNase R (Lucigen, RNR07250) at 37°C for 30 min and then placed on ice. Water was used in mock reactions. After incubating the sample with 1 μl of 1mM EDTA, 1 μl random hexamer (100 μM) and 1 μl of 10 mm dNTPs at 65 °C for 5 min, the denatured RNA was used for the cDNA synthesis with a Superscript IV RT system (ThermoFisher, 18091050) according to manufacturer’s instructions. For GFP qRT-PCR, total RNA was extracted from cells using the RNeasy Mini Kit, and the genomic DNA was extracted from cells using Quick Extract DNA Extraction Solution (ThermoFisher, NC9904870) according to the instructions. The total RNA was used for the cDNA synthesis with iScript cDNA Synthesis Kit (Biorad, 1708891) according to manufacturer’s instructions.End point PCR was performed with SapphireAmp (Takara, RR350B). Cycling conditions for circRNA were as follows: 95 °C 5 min, followed by 40 cycles of 95°C 1 min, 62°C 1 min, 72 °C 2 min, and a final elongation step at 72°C for 10 min. Cycling conditions for linear mRNA is the same condition with 25 cycles. For end point PCR, 2 μl cDNA templates were used for both circular and linear PCR reactions. For qRT-PCR, cDNA products and DNA products were diluted to 1:10 and 2 μL was used as template for real time PCR reaction with PowerUp SYBR Green (Applied Biosystems, A25779). The primers used for both end point PCR and qRT-PCR analysis are listed in Supplementary Table S1.

### Northern blots

Northern blotting for circular RNA analyses was performed as described previously (29). Briefly, total RNAs were extracted using the TRIzol reagent (Invitrogen). 20–30 μg of total RNA from cancer cell lines was treated with RNase R or 8 μg of total RNA used for mock treatment. RNA samples then separated on 1.5–2.0% formaldehyde agarose gels in MOPS buffer. The RNA was transferred to the Hybond-N+ membrane (GE Lifesciences) with 10xSSC. RNA hybridization was carried out in PerfectHyb buffer (Sigma) overnight at 65 °C. Probes were produced by PCR amplification in the presence of [α-^32^P] dCTP. Primers used for generating the probes are listed in Supplementary Table S1.

### Patients and tissue specimen collection

Frozen MCC tumors and normal skin were obtained from the UTSW Tissue Management Shared Resource. Tissue collection was performed with written informed consent from the patients and the studies were approved by the UT Southwestern institutional review board. Excess FFPE tissues were obtained from patients diagnosed with Trichodysplasia Spinulosa by clinical and histological criteria. The UT Southwestern Institutional Review Board approved these retrospective studies and the need for consent was exempted.

### Actinomycin D treatment

WaGa suspension cells were grown in six-well plates. cells were incubated with 2 μg/ml Actinomycin D (Sigma-Aldrich) or DMSO as a control and collected at indicated time points. 2 μg total RNA was subjected to 5 U RNase R treatment at 37 °C for 30 min. After the cDNA synthesis, the transcript level of circATLO and ALTO mRNA were analyzed by qRT-PCR. After 293T cells were grown overnight in six-well plates, cells were transfected with pCDNA-GFP reporter vector (1ug) and indicated plasmids including pCDNA control vector (1 ug), Flag-circALTO1 (1 ug) and Flag-circALTO2 (1 ug) plasmids. After 48 hours, cells were incubated with 2 μg/ml Actinomycin and collected at indicated time points.

### Nuclear and Cytoplasmic fractionation

Fractionation was performed according to the protocol performed before (29). Typically, cells grown on 35mm dishes were used as the starting material. After being trypsinized and pelleted, cell were resuspended in 250 μl of ice-cold Buffer I [0.5% Triton X-100, 0.32M sucrose, 3mM CaCl2, 2mM MgCl2, 0.1mM EDTA, 10mM Tris (pH 8.0), 50mM NaCl, 1mM DTT, 0.04 U/μl RNase inhibitor (ThermoFisher, 18091050)]. After a 15 min incubation on ice, cells were centrifuged at 500 g for 5 min at 4 °C. The supernatant was collected for the cytoplasmic fraction and the pellet was resuspended in 250 μl of Buffer I for the nuclear fraction. Afterward soluble fraction RNA was then extracted using RNeasy Mini Kit (Qiagen, 74104) and cDNA synthesis with Superscript IV RT system (ThermoFisher, 18091050) using random hexamers. The transcript levels were examined using gene specific primers by qRT-PCR.

### Luciferase reporter assay

The luciferase reporter assay performed as described before (10). In brief, 293 or 293T cells in 12-well plates were transfected with 5ng of the pCDNA3.1dsRlucMCVTAg plasmid, as well as 5ng of pCDNA3.1dsFFluc as a transfection control, along with 1 ug of the indicated plasmids per well (pCDNA-EV 1ug, pCDNA-EV 800ng + MCPyV miRNA 200ng, MCPyV miRNA 200ng + circALTO1 800ng, MCPyV miRNA 200ng + circALTO2 800ng). After 48 h transfection, cells were harvested and analyzed using a dual luciferase assay system (Dual-Glo luciferase assay system; catalog no. E2980) according to manufacturer’s instructions (Promega). Results are presented with the *Renilla* luciferase levels normalized by the firefly luciferase levels. To test for transcriptional activation, 293 cells in 12-well plates were transfected with 5 ng of the pCDNA3.1dsRlucMCVTAg and pCDNA3.1dsFFluc plasmid along with 1 ug of either pCDNA-EV, circALTO1, or ALTO1-SASD. Luciferase expression was analyzed 48 hr post-transfection.

### Constructs and oligonucleotide transfection

Wild type and mutant circALTO constructs were generated by gene synthesis (IDT or GeneWiz) based on the reference sequences for MCPyV (NC_010277.2) and TSPyV (NC_014361.1). Exact sequences listed in Supplementary Table S1. Constructs were PCR amplified (AccuPrime, ThermoFisher 12344040) and cloned into pCDNA3.1 after restriction digest with the appropriate enzymes. Constructs were confirmed by Sanger Sequencing. Constructs were transiently transfected into 293T cells with Lipofectamine 3000 reagents (ThermoFisher) according to manufacturer’s instructions. After 48–72 h transfection, cells were harvested for fractionation, RNA preparation, and WB. The pU-5864 reporter plasmid, a gift from Chris Buck, contains the EF1-*α* promoter and 5’ untranslated region in addition to the upstream regulatory region from BPV1 (40).

All siRNAs were synthesized by Sigma-Aldrich. The 293T cells co-transfected with 80 pmol of the indicated siRNA and 1 μg of the indicated constructs using Lipofectamine 3000 without P3000 reagent. After 68-72 h transfection, cells were harvested for RNA preparation, WB, and m6A IP. The oligos used in this study are listed in Supplementary Table S1.

### m^6^A RNA immunoprecipitation assay

Total RNA was harvested 48–72 h post-transfection and extracted by (Qiagen, 74104). 5 μg total RNA was incubated with 1 μg m6A polyclonal antibody (Synaptic Systems# 202003)(41) and beads with rotation at 4°C. Immunoprecipitated RNA was then purified using QIAgen RNeasy columns. Purified immunoprecipitated RNA, along with 10% input RNA were then reverse transcribed by Superscript IV RT system (ThermoFisher, 18091050) using random hexamers. The modification levels were examined using gene specific primers by qRT-PCR. Percentages of m6A modified RNA for both circALTO and SON were calculated based on the input amount.

### Western blotting

Whole cells were trypsinized and harvest subjected to Laemmli sampling buffer and boiled for at 95 °C for 10 min. The protein extracts were separated on SDS-PAGE gels, transferred to PVDF membranes and incubated with indicated antibodies. The antibodies were purchased from commercial sources: anti-FLAG HRP (1:1000, Sigma, A8592), anti-HSP90 (1:1000, Cell Signaling Technologies, 4877S), anti-CD63 (1:600, BD Biosciences, 556019), anti-CD81 (1:600, R&D Systems, MAB4615), anti-Calnexin (1:1000, Cell Signaling Technologies, 2679T), anti-*β*-Tublin (1:1000, Proteintech, 10094-1-AP), anti-Mettl3 (1:1000, Proteintech, 15073-1-AP), anti-GFP (1:500, Santa Cruz Biotechnology, sc-9996) and TSPyV pAbMT (1:100) (13). Membranes were then incubated with the appropriate HRP-conjugated secondary antibody (1:000, anti-mouse IgG, Cell Signaling Technologies, 7076P2; 1:1000, anti-rabbit IgG, ThermoFisher, G21234) and developed with an ECL system (Perkin Elmer, NEL104001).

### Immunofluorescence

The 293 T cells were plated into Nunc 4-well chamber slides (ThermoFisher 154453) and transfected with indicated plasmids and siRNA. After 48 h transfection, cells were fixed in 4% paraformaldehyde at RT for 10 min, followed by 0.1% Triton X-100 permeabilization at RT for 10 min. Cells were then incubated with FLAG antibody (1:100, Sigma, F1804) overnight, then stained with Alexa 488-conjugated secondary antibody (ThermoFisher). Samples were then subjected to DAPI nuclear counterstain (Vector Labs), and imaged by fluorescent microscope.

### Total exosome isolation and transfer

Exosome isolation was performed according to a published protocol (42). Cell-conditioned medium was collected grown for 3 days in cell culture flasks (WaGa cells) or 150 mm plates (MCC26, pCDNA and Flag-circALTO1/2 transfected 293T cells) with exosome-depleted FBS (Invitrogen). The collected media was first centrifuged at 300×g for 10 min at room temperature to pellet and remove cells. The following centrifugation steps were performed at 4 °C. In brief, the supernatant was spun at 2,000×g for 15 min. Then, the supernatant was centrifuged at 10,000×g for 60 min. Next, the media supernatant was passed through a 0.22 µm pore PES filter (Millipore). This supernatant was ultracentrifuged at 100,000×g for 90 min in a SW-28Ti Rotor Swinging Bucket rotor (Beckman Coulter) and resuspended in PBS, then centrifuged at 100,000×g for 90 min again. The resulting Exosomes pellet resuspended in 250 µL of PBS and stored at −80 °C for further use. Exosomes were then subjected to RNA isolation, western blotting and TEM. The RNA isolated form exosome using Total Exosome RNA & Protein Isolation Kit (Invitrogen). For western blotting, the exosomes in PBS directly subjected to Laemmli sampling buffer with 2-mercaptoethanol or without 2-mercaptoethanol (CD63 and CD81) and boiled at 95 °C for 10 min.

### Transmission electron microscopy (TEM)

Exosome samples were adsorbed to glow-discharged carbon-coated 400-mesh copper grids for 3 minutes and then stained with 2% (weight/volume) uranyl acetate in water, rinsed briefly with water and air dried. The grids were visualized on a FEI Tecnai 12 TEM at 80 kV. Images were captured on a Gatan Orius CCD camera with Gatan Digital Micrograph software.

### Nanoparticle tracking analysis (NTA)

Samples were diluted 1:100 for nanoparticle tracking analysis with the ZetaVew particle analyzer (Particle Metrix). The size (nm) and concentration (cm^-3^) was used to determine the extracellular vesicle size using GraphPad Prism.

### Transfer and DiI-Labeled Exosomes to HeLa cells

HeLa cell were planted in 24 well plates and grew until 70% confluence to use. Cells washed twice with PBS and incubated with 100 μl indicated exosomes for 70 hours at 37 °C in exosomal-depleted FBS (Invitrogen) supplemented media. Whole cells were washed twice with PBS, trypsinized and harvest subjected to Laemmli sampling buffer with 2-mercaptoethanol and boiled for at 95 °C for 10 min. 1 μM DiI (Invitrogen) labeled with purified exosomes as previously described (43). Unbound DiI were removed by Exosome Spin Columns (Invitrogen). Purified exosomes were added to HeLa cells and incubated for 24 h. Recipient cells were then washed in PBS, fixed in 4% paraformaldehyde, and imaged by fluorescent microscope. For GFP transcript assay, 100 μl purified exosomes were added to HeLa cells in 24-well plates and incubated for 24 h, then cells were transfected with pCDNA-GFP reporter vector. After 48 hours, cells were collected for experiments.

## RESULTS

### Identification of Merkel cell polyomavirus circRNAs

To determine whether MCPyV might generate circRNA, we used the previously described vircircRNA pipeline on RNA-Seq datasets of MCPyV infected cells (https://github.com/jiwoongbio/vircircRNA) (29). Three potential circRNAs in the early region were predicted to share the same 5’ splice site, but distinct 3’ splice site, and all three encompassed both the previously described MCPyV-encoded miRNA docking sites (complementary to the miRNA locus) and the ALTO/MT ORF (Figure 1A, S1A). Additional potential circRNAs were also identified upstream of the VP2 minor capsid protein ORF in a region where some polyomaviruses encode an agnoprotein gene.

**Figure 1.**
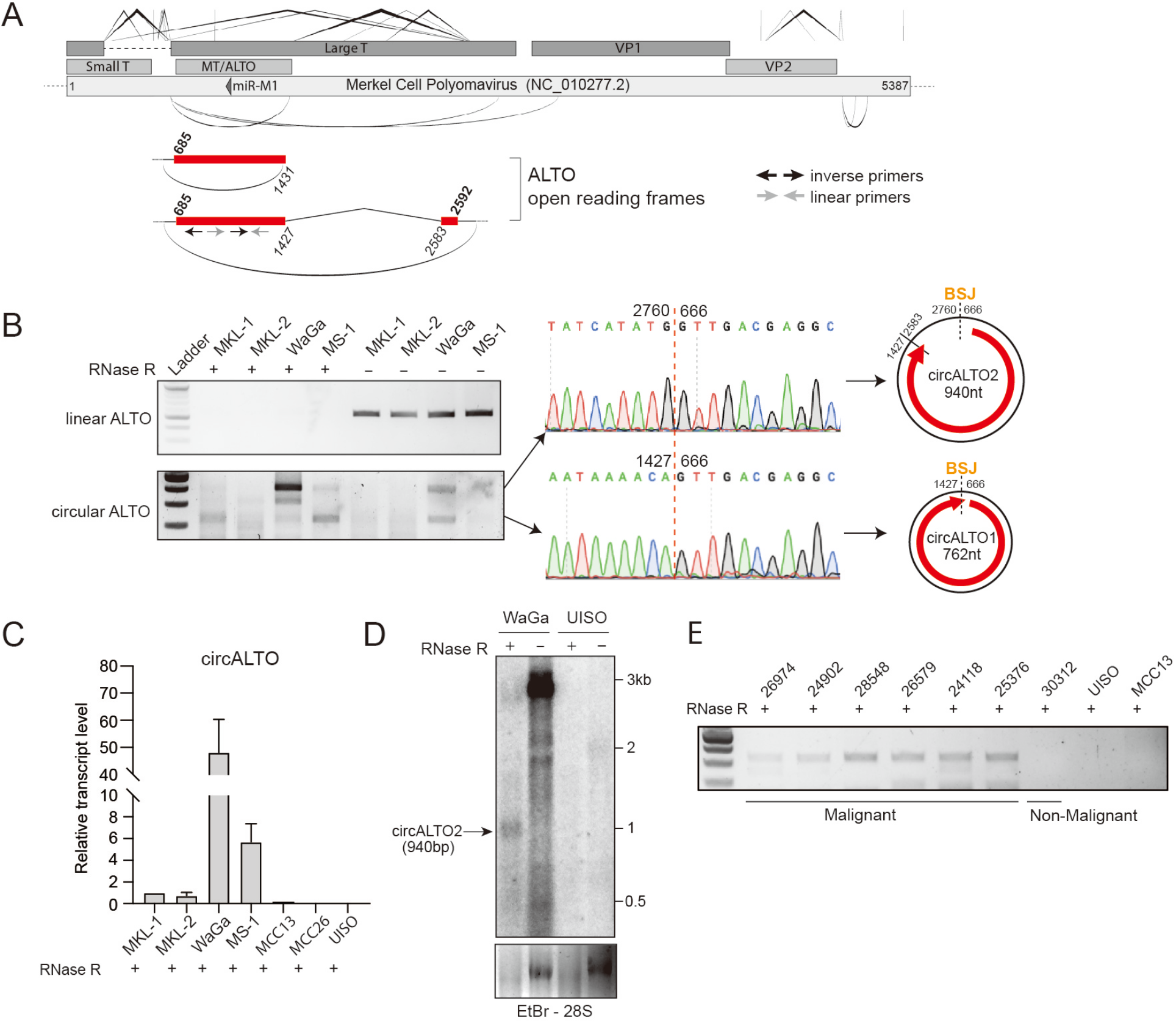
Identification of Merkel cell polyomavirus circRNAs. (A) (Top panel) Schematic representation of potential circRNAs identified by the vircircRNA for MCPyV. V-shaped lines above the map indicate forward splicing events. Elliptical arcs below the map indicate predicted backsplicing. (Bottom panel) Red rectangles indicate ALTO open reading frames. Bold numbers above the map indicate the first base of ALTO start and stop codons, italic numbers below the map indicate the positions of exon boundaries. (B) (Left panel) RT-PCR analysis of four VP-MCC cell lines (MKL1, MKL-2, MS-1 and WaGa) with and without RNase R treatment. (Middle panel) Sanger sequencing of inverse PCR products from WaGa cells confirmed the backsplice junction of the predicted circRNAs. (Right panel) Schematic of the two different circALTO forms, we termed circALTO1 and circALTO2 based on the product length. (C) qRT-PCR analysis for circALTOs from VP-MCC (MKL-1, MKL-2, MS1, and WaGa) and VN-MCC (MCC13, MCC26 and UISO) after RNase R treatment. *ACTB* served as the normalization control. Values are the mean of three technical replicates; bars represent standard deviation. (D) Northern blot of total RNA after with and without RNase R treatment from VP-MCC cell line WaGa and VN-MCC cell line UISO using an ALTO-specific probe. Arrow indicates RNase R resistant band consistent with circALTO2. (E) Endpoint PCR confirmed the presence of circALTO2 in patient MCC tumors, but not a non-malignant skin control or VN-MCC cell lines.

To confirm the presence of circRNA transcripts, we used inverse PCR on four VP-MCC cell lines, MKL-1, MKL-2, MS-1, and WaGa. Inverse PCR products could be identified for the predicted circRNAs in the early region, but not adjacent to the late region, likely because the expression of late genes is typically down-regulated in MCC tumors (Figure 1B, data not shown). We chose to focus on the characterization of the potential circRNAs in the early region (circALTO).

RNase R, an exonuclease that degrades most linear ssRNAs, was used to distinguish between linear RNAs arising from multiple passes around the viral genome and RNAs circularized by backsplicing. Total RNA from four VP-MCC cell lines was subjected to inverse RT-PCR targeting circALTO. PCR products remained detectable after RNase R digestion, indicating the presence of circular RNAs (Figure 1B). The RNase R resistant inverse PCR products from WaGa cells were then sequenced. The slower-migrating band, circALTO2, represents a 940nt circRNA with a ‘backspliced’ ALTO ORF that encompasses an additional canonical splice that has previously been described for linear ALTO mRNAs. The faster-migrating, circALTO1, is a 762nt circRNA containing most of the ALTO ORF (Figure 1B). Other intermediate-sized inverse RT-PCR bands observed in VP-MCC cells were found to contain insertions not derived from the MCPyV genome (Supplementary Figure S1B). Next, we compared levels of circular ALTO RNAs from RNA prepared from VP-MCC and virus-negative (VN)-MCC lines (MCC13, MCC26, and UISO) with RNase R treatment. We designed the PCR primers in a region shared by both circALTO1 and circALTO2. Both endpoint and qPCR revealed that circALTO could be detected in VP-MCC, but not VN-MCC cells (Figure 1C, S1C). The presence of circALTO2 in RNase R-treated total RNA preparations of WaGa cells was confirmed by northern blotting (Figure 1D). Finally, we next tested for the expression of circALTO in patient tumors (Figure 1E, Supplementary Figure S1D-E). Endpoint inverse RT-PCR products representing circALTO2 were detected after RNase R-digestion for 6/7 VP-MCC tumors, but not in non-malignant skin controls or VN-MCC lines (Figure 1E). Despite its detection in VP-MCC cell lines, circALTO1 could not be detected in the tumor tissues suggesting that the abundance of the circALTO isoforms might differ in vivo.

### Characterization of circALTO

Circular RNAs have been reported to be more stable than linear RNA. Indeed, after treatment with the transcriptional inhibitor, Actinomycin D (30), a time course qPCR analysis showed that circALTOs were significantly more stable than linear ALTO mRNAs, with a half-life exceeding 24 hours in WaGa cells (Figure 2A). Next, we investigated the localization of circALTO in WaGa cells. Cellular fractionation analyses showed that circALTOs mainly localized to cytoplasm while the majority of linear ALTO RNA localized to the nucleus (Figure 2B). Thus, circALTOs are very stable and largely localized to the cytoplasm. Because N^6^-methyladenosine (m^6^A) modifications have been reported to be abundant on circRNAs and have been reported to promote m^6^A reader protein binding, splicing, and cap-independent translation (44-46), we performed m^6^A RNA immunoprecipitation to determine whether the circALTO RNAs might be modified. In WaGa cells, we found both SON, a positive control mRNA, and circALTO were pulled down by m^6^A antibody precipitation, but not an IgG control antibody (Figure 2C).

**Figure 2.**
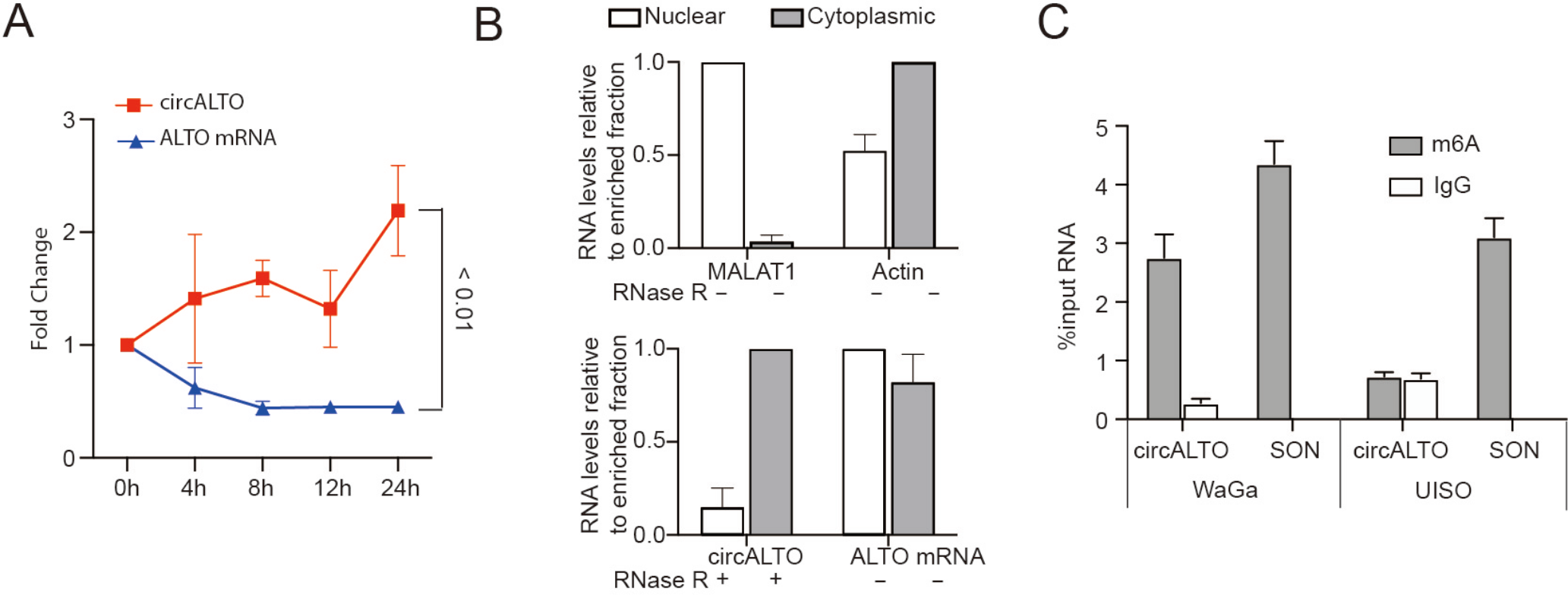
Characterization of circALTO. (A) The expression of circALTOs and its linear mRNA counterparts after treatment with Actinomycin D at the indicated time points in WaGa cells by qRT–PCR analysis. Both circALTO and linear ALTO were normalized to *ACTB* and then compared to levels at the pre-treatment (0 h) time point. Error bars calculated the SD from 3 biological replicates. The *P* value was determined using two-tailed t-test. (B) WaGa were fractionated and analyzed by qRT-PCR. MALAT1 and *ACTB* served as nuclear and cytoplasmic fractionation controls respectively. Values normalized to the enriched fraction. Error bars were calculated from 3 biological replicates. (C) qRT-PCR of m^6^A or IgG control RNA immunoprecipitation (RIP) of VP-MCC cell lines WaGa and VN-MCC UISO. SON, m^6^A RNA IP control. Error bars represent SD (n = 3 biological replicates).

### Confirmation using recombinant circALTO expression vectors

To evaluate potential functions of circALTO, we generated expression vectors encompassing the entire backspliced region of ALTO gene (Supplementary Figure S2A). All constructs included Quaking (QKI) protein-binding sites which have been previously been shown to enhance the circularization of RNAs (47) and an amino(N)-terminal 3xFlag epitope tag (Supplementary Figure S2A). The circALTO2 construct contains an additional exon which included an in-frame stop codon. In contrast, circALTO1 does not naturally contain an in-frame stop codon when circularized, and thus appears to encode for a circular ORF capable of ‘infinite’ rolling circle translation (Supplementary Figure S2A). The circALTO constructs were transfected into HEK293T cells, and RNase R-resistant circRNAs were detected by RT-PCR from both WT and epitope-tagged circALTO (Supplementary Figure S2B, left). Sanger sequencing of the RT products confirmed that the circALTO construct generated the expected backsplice junctions (Supplementary Figure S2B, right). As with the circular MCPyV genome and integrated tandem repeats of the viral genome, read-through transcription of the entire pCDNA3.1-based plasmid followed by forward splicing between distinct copies of the ALTO region could theoretically generate “ false backsplice” junctions. Indeed, primers amplifying the CMV promoter region could be detected by RT-PCR, indicating that readthrough transcription of the pCDNA vector could occur. However, readthrough transcription products could no longer be detected after RNase R treatment while the relative detection of circALTO backsplice junctions increased after RNase R treatment (Fig. 1B, Supplementary Figure S2C). Finally, northern blots confirmed that an RNase R resistant RNA of the expected size of circALTO2 was generated by the circALTO2 construct (Supplementary Figure S2D). CircALTO1 could not be reliably distinguished above background signals, possibly due to its smaller size. Fractionation and quantitation of nuclear and cytoplasmic circALTO from 293T cells transfected with circALTO1 (Supplementary Figure S2E) demonstrated that circALTO1 was enriched in the cytoplasmic fraction, consistent with the findings from WaGa cells (Figure 2B).

### CircALTOs do not sponge MCPyV miR-M1

Many circRNAs function as microRNA (miRNA) sponges (31,48,49), and the MCPyV ALTO locus encodes for miRNAs, miR-M1-3p and miR-M1-5p are complementary to the circALTO transcript (10,11). Therefore, we tested whether circALTOs might bind and inhibit the MCPyV miRNA. We used the circALTO constructs together with a luciferase reporter construct to investigate the ability of circALTOs to bind to MCPyV miRNA in transfected cells. A reporter plasmid that contains Renilla luciferase fused to a 300 nucleotide region of the MCPyV early transcript shows decreased luciferase expression when co-transfected with an MCPyV miR-M1 (MCC350 miRNA variant) expressing plasmid (10). Using these plasmids, we tested whether circALTO might be able to “ sponge” miR-M1-5p/3p and thereby rescue miR-M1 mediated silencing of the early region. When we co-transfected a MCPyV miRNA-M1 expression vector and a circALTO1/2 expression vector, luciferase expression was not rescued compared to the HEK293 cells transfected with MCPyV miRNA alone (Figure 3A). This inability to rescue luciferase expression was confirmed in HEK293T cells, and was also observed when we assayed a sequence variant MCPyV miRNA (MCC339) (Supplementary Figure S3A-B). These results suggest that the circALTO does not function as an inhibitor for MCPyV miR-M1.

**Figure 3.**
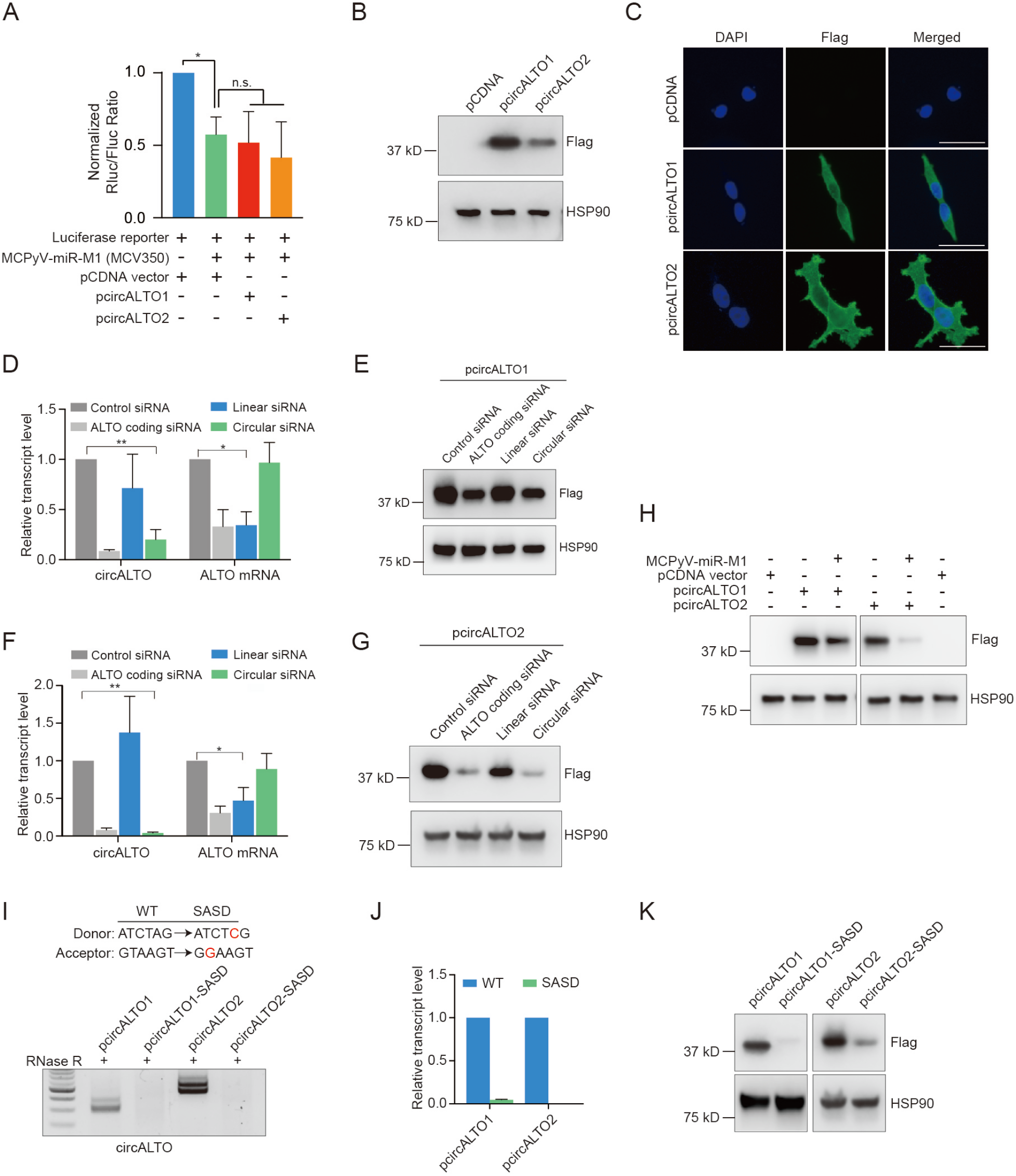
MCPyV circALTO can be translated. (A) CircALTOs do not inhibit MCPyV miRNA function in vitro. 293 cells were transfected with *Renilla* luciferase reporter with a 300 nucleotide region complementary to the MCPyV miRNA and the indicated plasmids including a MCPyV-miRNA-M1 expression plasmid (derived from isolate MCV350), pCDNA3.1 control vector, pCDNA circALTO1 or pCDNA circALTO2 expression plasmid. Firefly luciferase served as a transfection control and *Renilla* luciferase levels are plotted normalized relative to firefly luciferase levels (n=2 biological replicates). The *P* value was determined by unpaired two-tailed t-test. (B) Western blot for FLAG from 293T cells co-transfected with a pCDNA vector control plasmid or pCDNA3.1-Flag-circALTO1/2. HSP90, loading control. (C) 293T cells were transfected with vector, the Flag-circALTO1/2 plasmids. After 48 hours, cells were fixed and stained for FLAG (green) and DAPI (blue). Scale bar = 50 μm. (D) qRT-PCR analysis for circALTO and ALTO mRNA in co-transfected Flag-circALTO1 constructs 293T cells treated with the indicated siRNAs (see Supplementary Figure S3A). *ACTB* served as the internal control. Error bars represent the SD (n = 3 biological replicates). (E) Western blot for FLAG from 293T cells co-transfected with Flag-circALTO1 and the indicated siRNA. HSP90 serves as loading control. (F) qRT–PCR analysis for circALTO and ALTO mRNA in co-transfected Flag-circALTO2 constructs 293 T cells with siRNAs (see Supplementary Figure S3A). *ACTB* served as the internal control. Error bars represent the SD (n = 3 biological replicates). (G) Western blot for FLAG from 293 T cells co-transfected with Flag-circALTO2. HSP90 is the loading control. (H) Immunoblotting analysis of 293T cells were transfected with vector, pCDNA-circALTO1, or pCDNA-circALTO2 with or without MCPyV miRNA expression vector. Co-transfection with the MCPyV miRNA expression vector results in decreased expression of ALTO. (I) Endpoint RT-PCR analysis for circALTO from 293T cells transfected with Flag-circALTO1/2 (WT) and the indicated constructs harboring splice sites mutations (SASD) (see Supplementary Figure S3A). (J) qRT-PCR analysis for circALTO from cells with WT constructs and constructs harboring SASD mutant showed (see Supplementary Figure S3A). *ACTB* served as the internal control. Error bar representative of 3 independent experiments. (K) Western blots for Flag circALTO from cells with WT constructs and constructs harboring the SASD mutation. HSP90, loading control. The *P* value was determined by unpaired two-tailed t-test, *<0.05, **<0.005.

### CircALTOs can be translated

We next investigated whether circALTOs might be translated. Transfection of Flag epitope-tagged circALTO expression vectors in 293T cells resulted in the production of proteins whose size was consistent with the previously described MCPyV ALTO protein (Figure 3B). Immunofluorescence assays showed that cells transfected with circALTO1/2 generated Flag-ALTO proteins which were localized in the cytoplasm (Figure 3C). To confirm that ALTO was encoded by the circular, rather than linear, ALTO RNAs, we used small interfering RNAs (siRNAs) to target the expression of ALTO RNA isoforms in circALTO1/2 transfected cells. Several siRNAs were designed for both circALTO1 and circALTO2 constructs: one targeting the circALTO1 or 2 backsplice junction (BSJ), a second targeting a linear region outside the backsplice exon, and a final siRNA targeting sequences shared by both the circular and linear RNAs within the ALTO ORF coding region. The efficiency of siRNA knockdown was confirmed by qRT-PCR analysis; siRNAs against the BSJ sequences specifically knocked down circALTO1/2, the siRNA against the linear sequences only significantly knocked down the linear ALTO transcript, and the siRNA targeting the ALTO coding sequences significantly decreased both circular and linear ALTO transcript levels (Figure 3D, F). Notably, the BSJ- and ALTO-targeting siRNAs strongly downregulated ALTO expression, whereas specific knockdown of the linear transcript only modestly impacted ALTO expression levels (Figure 3E, G). In addition, in immunofluorescence experiments, cells co-transfected circALTO1/2 plasmids and BSJ specific siRNAs showed marked reductions in Flag staining compared to circALTO1/2 control transfections (Supplementary Figure S3C). The results indicate that a majority of ALTO protein expression in this system is attributable to circALTO RNA, particularly in the case of the ALTO2 isoform.

The annealing of fully complementary miRNA’s can result in the Ago2-dependent cleavage and degradation of mRNA (50). While circALTO1/2 did not appear to possess an ability to sponge miR-M1, we next tested whether circALTO1/2 stability might be impacted by miR-M1-5p/3p binding. We co-transfected the circALTO1/2 and miR-M1 expression plasmids and found that ALTO expression levels were markedly decreased compared to cells transfected with circALTO1/2 alone (Figure 3H). Thus, rather than sponging miR-M1, circALTO is negatively regulated by miR-M1.

N^6^-methyladenosine (m^6^A) modifications have been implicated as critical modifications in circRNA expression (44). To assess the role of m^6^A modification in circALTO expression, we used siRNA to knock down Mettl3, a conserved m^6^A methyltransferase. While Mettl3 knockdown somewhat reduced the amount of circALTO RNA pulled down by anti-m^6^A immunoprecipitation, levels of ALTO protein did not show marked reductions after Mettl3 siRNA treatment (Supplementary Figure S3D-G). Finally, to confirm that backsplicing was necessary for the expression and translation of circALTO1/2, we generated circALTO constructs with mutations in the splice acceptor and splice donor sites (SASD) (Figure 3I). As expected, both endpoint and qRT-PCR (Figure 3I-J) analyses did no show detectable levels of circALTO transcripts from the constructs with splice site mutations. Consistent with a critical role of the circALTO isoforms in translation, ALTO protein levels were also markedly downregulated in the splice site mutant constructs compared with wild type control (Figure 3K). In summary, we find that circALTO1 and circALTO2 generate ALTO proteins in cultured cells.

### circALTOs increase expression from the human cytomegalovirus (CMV) immediate early promoter

While MCPyV LT and sT antigens play critical roles in viral replication and tumorigenesis, the biological functions of MCPyV ALTO remain unknown. While testing circALTO in miRNA-interference assays, we noticed that co-transfection of either pcircALTO1 or pcircALTO2 appeared to increase the luciferase expression from reporter plasmids encoding Renilla luciferase (RLuc) or firefly luciferase (FFLuc) under control of the immediate early promoter (Figure 4A). Increased expression of co-transfected luciferase plasmids required circularization of the ALTO1 RNA, as luciferase expression was not significantly increased when the co-transfected with pcircALTO1 plasmid contained mutations in the backsplice acceptor and donor sites (circALTO-SASD) (Figure 4A). To verify this observation, we tested a CMV promoter-driven green fluorescent protein plasmid, pCDNA-GFP, and confirmed that that the co-transfection of either pcircALTO1 or pcircALTO2 resulted in markedly increased levels of GFP expression by both Western blot and immunofluorescence (Figure 4B-C). GFP transcript levels were similarly increased, as qRT-PCR revealed that pcircALTO co-transfection resulted in a ∼4-fold increase in GFP transcript levels (Figure 4D). Increased expression of co-transfected luciferase plasmids required ALTO circularization, as luciferase expression was not significantly increased when the co-transfected with circALTO plasmid contained mutations in the backsplice acceptor and donor sites (circALTO-SASD) (Figure 4A). As the circALTO-SASD inhibits both circularization and ALTO expression, we next tested whether ALTO expression was required for the upregulation of co-transfected reporter plasmids. Thus, we generated a mutant circALTO in which all potential ATG start codons had been mutated (circALTO1-*Δ*ATG) (Figure 4E). While circALTO1-noATG could still efficiently circularize (Figure 4E), its ability to encode for ALTO protein was abrogated (Figure 4F). Notably, the circALTO1-*Δ*ATG mutant was no longer able to increase the expression of GFP when co-transfected with the pCDNA-GFP reporter plasmid as assessed by both Western and qRT-PCR (Figure 4F-G).

**Figure 4.**
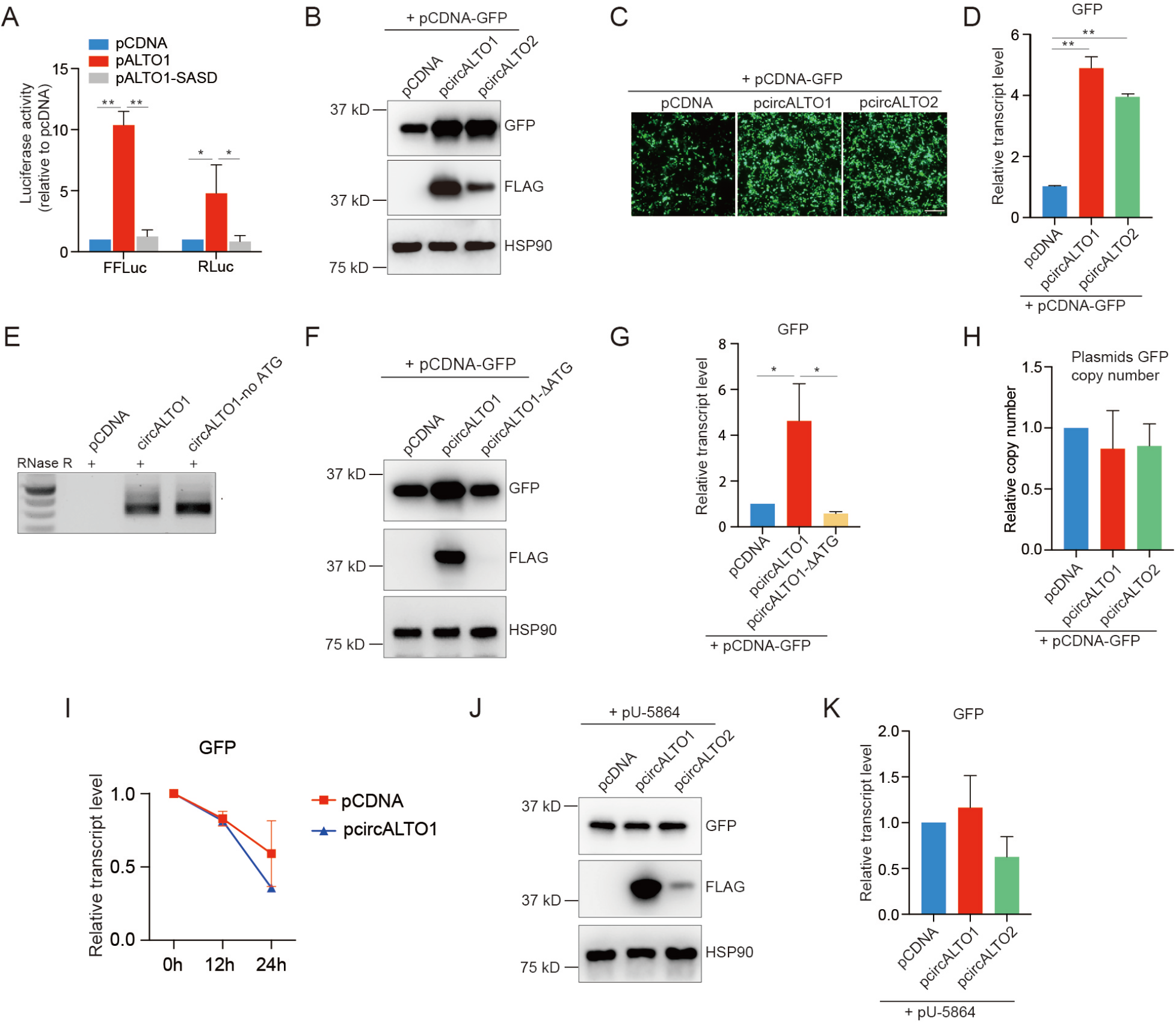
MCPyV circALTO increases transcription by the CMV promoter. (A) 293 cells were co-transfected with *Renilla* luciferase reporter or Firefly luciferase reporter and with either pCDNA empty control, circALTO1, or circALTO1-SASD (n=3 biological replicates). The *P* value was determined by unpaired two-tailed t-test. (B) Western blot for GFP and FLAG from 293T cells which were co-transfected with pCDNA-GFP and indicated plasmids including pCDNA control vector, Flag-circALTO1, or Flag-circALTO2 plasmids. HSP90 is the loading control. (C) 293T cells were co-transfected with pCDNA-GFP vector and indicated plasmids. IF images are representative of GFP expression after 48 hours. Scale bar = 200 μm. (D) qRT-PCR analysis for GFP from cells with pCDNA-GFP constructs and indicated plasmids. *ACTB* served as the internal control. Error bar representative of 3 independent experiments. (E) A construct in which all potential ATG start codons have been mutated, circALTO1-noATG, was generated. RT-PCR analysis for circALTO of total RNA prepared from 293T cells transfected with pCDNA, circALTO1, and circALTO1-noATG indicate that both circALTO1 constructs are efficiently circularized. (F) Western blot for GFP and FLAG from 293T cells were transfected with pCDNA-GFP reporter vector and pCDNA control vector, Flag-circALTO1, and Flag-circALTO1-noATG plasmids. HSP90 is the loading control. (G) qRT-PCR analysis for GFP from 293T cells are shown as in (F). *ACTB* served as the internal control. Error bars = SD from one biological replicate. Results are representative of 3 independent experiments. (H) 293T cells were co-transfected with pCDNA-GFP plasmid and either pCDNA empty control, circALTO1, or circALTO2. Levels of pCDNA-GFP detected from DNA were assessed by qPCR. (I) 293T cells were co-transfected with pCDNA-GFP and pCDNA control vector or circALTO1. After 48 hours, actinomycin D added to the tranfected cells. The expression of GFP after treatment with Actinomycin D at the indicated time points by qRT–PCR analysis. Error bar = SD from one biological replicate. Results are representative of 2 independent experiments. GFP were first normalized to 18S and then normalized to levels at the pre-treatment (0 h) time point. (J) Western blot for GFP from 293T cells were co-transfected with pU-5864 and either pCDNA control, circALTO1, or circALTO2 plasmids. HSP90 serves as the loading control. (K) qRT-PCR for GFP from cells co-transfected with pU-5864 constructs and indicated plasmids. *ACTB* served as the loading control. Error bar = SD from one biological replicate. Results are representative of 3 independent experiments. The *P* value was determined by unpaired two-tailed t-test, *<0.05, **<0.005.

To determine how ALTO might increase GFP transcript levels, we first determined whether reporter plasmid replication might be increased in 293T cells co-transfected with circALTO. However, copy number of pCDNA-GFP transcripts as assessed by RT-qPCR were not significantly increased after co-transfection with pCDNA control or circALTO1/2 expression vectors (Figure 4H). Next, we determined whether ALTO might alter the stability of reporter RNAs. After co-transfecting cells with the GFP reporter plasmid and vector or circALTO expression vector, cellular transcription was halted with Actinomycin D, and the stability of the GFP transcript was assessed by qPCR. Co-transfection of circALTO did not significantly alter the abundance of GFP RNAs 12 hours after transcription was halted, and GFP RNA stability in circALTO transfected cells was instead somewhat lower after 24 hours of Actinomycin D compared to the pCDNA transfected control (Figure 4I). Thus, ALTO does not appear to markedly increase GFP RNA stability, suggesting that ALTO might increase GFP transcription. The pCDNA3.1 vector utilizes the CMV promoter to drive transgene (GFP reporter) expression. To test whether ALTO might promote transcription in a promoter-specific fashion, we utilized a distinct reporter plasmid, pU-5864, in which GFP expression is driven by the human EF1-*α* promoter. Notably, circALTO co-transfection no longer increased GPF expression as assessed by WB or qRT-PCR when pU-5864 was used as the reporter plasmid (Figure 4J-K). We conclude that MCPyV ALTO appears to promote the expression of co-transfected genes by functioning as a transcriptional activator for specific promoters.

### Characterization of exosomes from circALTO expressing cells

Exosomes are 40-150 nm, membrane bound extracellular vesicles (EVs) that are enriched for specific proteins, lipids, and nucleic acids, including RNA. CircRNAs have been shown to be enriched in extracellular vesicles, including exosomes (35), and extracellular vesicles may play a role in the life cycle of polyomaviruses (51,52). Therefore, we investigated whether circALTOs could be detected in exosomes. EVs consistent with exosomes were isolated from a VP-MCC cell line, WaGa, and a VN-MCC cell line, MCC26 through differential ultracentrifugation. To confirm the purity of the EV isolation, purified EVs were subjected to immunoblot analysis to identify the presence of common exosome associated protein markers, including CD63 and CD81. EVs isolated from WaGa and MCC26 both expressed classical exosomal markers including CD63 and CD81, but did not show evidence of contamination by cytoplasmic proteins (*β*-tubulin) or intracellular membranes (calnexin), which were readily detected in whole cell lysates (Figure 5A). Purified EVs were also analyzed by nanoparticle tracking analysis (NTA). NTA of EVs isolated from WaGa and MCC26 cells showed a size distribution expected for exosomes (Figure 5B, Supplementary Figure S4A) (53). Finally, negative stain transmission electron microscopy (TEM) of WaGa EVs showed round and cup-shaped vesicles ranging from 50-150nm in size (Figure 5C).

**Figure 5.**
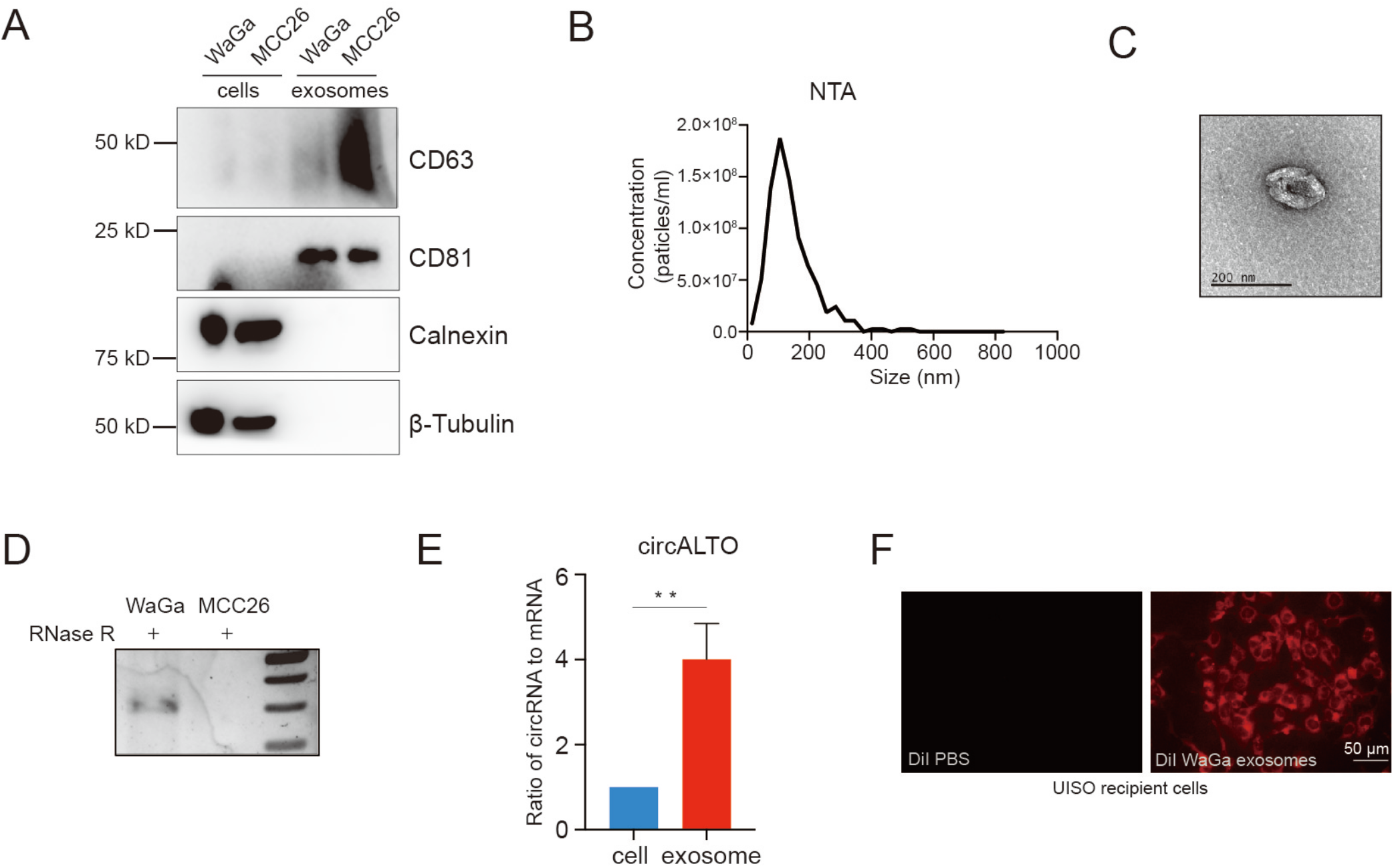
Characterization of exosomes from WaGa cells. (A) Western blot analysis of exosomal marker proteins in extracellular vesicles isolated from VP-MCC cell line WaGa and VN-MCC cell lines MCC26. CD63 and CD81 are exosomal marker proteins. The absence of calnexin and *β*-tubulin exclude contamination from the cytoplasm and cellular organelles. (B) Nanoparticle tracking analysis of extracellular vesicles isolated from WaGa cells are consistent with exosomes. (C) Representative TEM images of exosomes isolated from WaGa cells show the expected size and cup-shaped morphology described for exosomes. (D) RT-PCR analysis of RNase R treated total RNA prepared from exosomes of WaGa and MCC26 cells reveals the presence of circALTO in the exosomes. (E) qRT-PCR analysis the ratio of circRNA to mRNA between WaGa cells and cell-derived exosomes. Both circALTO and linear ALTO were first normalized to *ACTB*, then the ratio of circALTO:linear ALTO was set as 1. Error bars represent the SD (n = 3 biological replicates). Two-tailed t-test was performed, **<0.005. (F) Fluorescence of UISO cells incubated with DiI labeled exosomes from WaGa or Dil labeled PBS control by fluorescent microscopy. Scale bar = 50 μm.

Endpoint RT-PCR from RNA prepared from WaGa and MCC26 EVs revealed that circALTO2 could be detected in WaGa, but not MCC26 EVs (Figure 5D). Because circRNAs have been shown to be enriched in exosomes relative to their levels in the cell, we performed qRT-PCR analysis to measure the ratio of circular to linear ALTO RNA from WaGa cells. Indeed, there was a four-fold enrichment of circALTO in exosomes compared to its levels in total cellular RNA (Figure 5E).

Studies have demonstrated that exosomal mRNAs and ncRNAs, including EBV derived miRNAs, can be transferred between cells (54,55). The purified EVs were used to determine whether circALTO can be taken up by UISO cells through exosomes. UISO cells were cultured with WaGa exosomes. To do this, EVs were purified, fluorescently labeled with 1,1′-dioctadecyl-3,3,3′3′-tetramethylindocarbocyanine perchlorate (DiI), and incubated with UISO cells for 24 h to confirm the efficiency of uptake. Fluorescence was only detected in UISO cells incubated with labeled WaGa EVs, but not with PBS containing dye (Figure 5F), suggesting specific uptake of the EVs into endocytic compartments.

To test whether exosomal circALTO could be transferred between cells, we first tested whether circALTO could be purified from circALTO transfected 293T cells. Indeed, EVs containing circALTOs could also be detected in 293T cells transfected with pcircALTO1 or pcircALTO2 expression constructs. The purity of EVs isolated from 293T cells was also confirmed by western blot, NTA, and TEM (Figure 6A-C, Supplementary Figure S4B). Both circALTO1 and 2 RNAs could readily be detected in purified EVs from 293T cells (Figure 6D).

**Figure 6.**
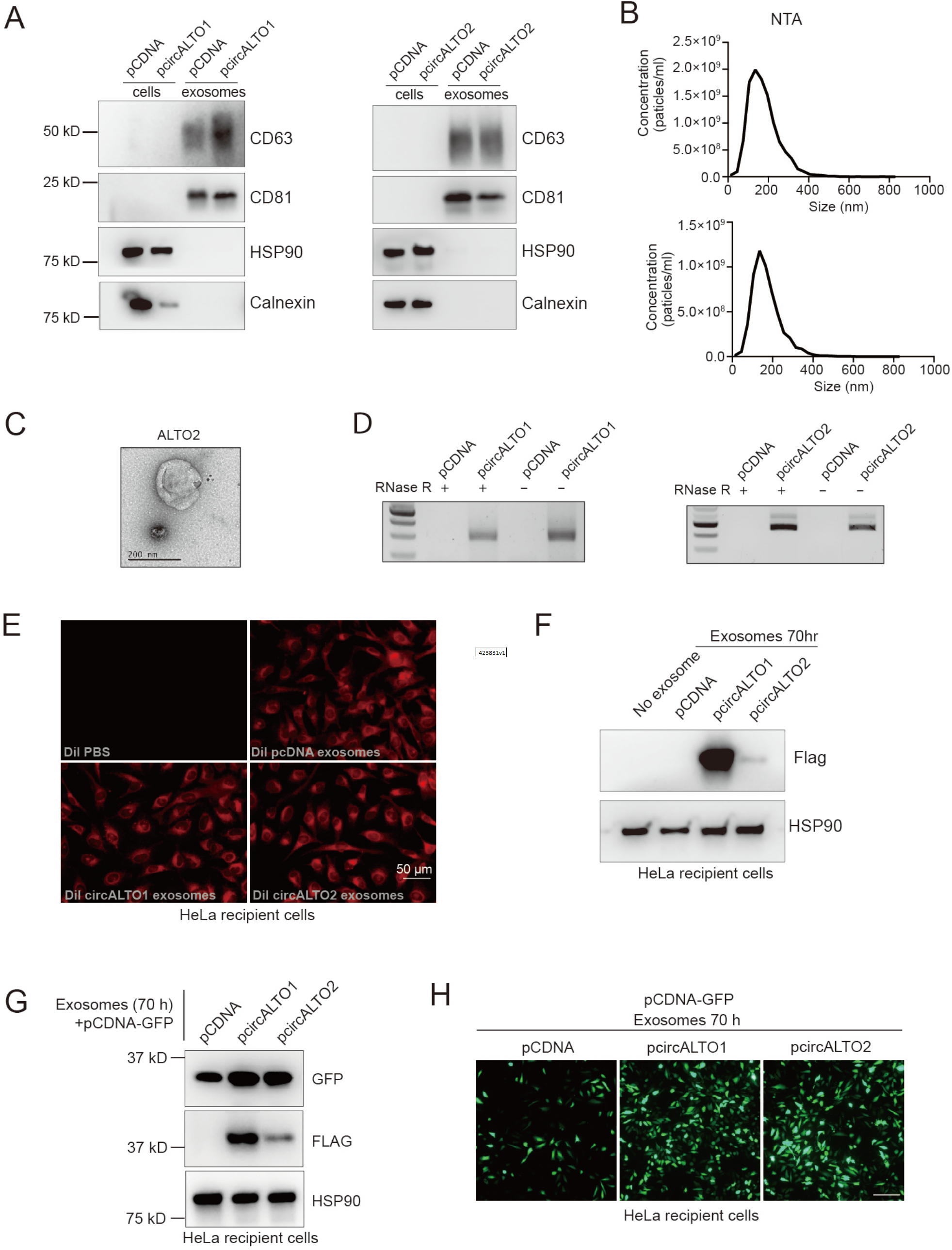
Exosomal circALTO can mediate ALTO expression in HeLa cells. (A) Western blot analysis of exosomal marker in exosomes extracted from 293 T cells transfected with pCDNA vector control or Flag-circALTO1/2. CD63 and CD81 are exosomal marker proteins, while calnexin and *β*-tubulin exclude contamination from the cytoplasm and cellular organelles. (B) Size distribution analysis of total exosomes isolated from the Flag-circALTO1 (upper panel) and Flag-circALTO2 (bottom panel) expressed 293T cells. (C) TEM images of exosomes isolated from circALTO2 expressed 293 T cells confirm expected cup-shaped morphology. (D) RT-PCR analysis of total RNA from exosomes of transfected with pCDNA vector control or Flag-circALTO1/2 293 T cells with and without RNase R treatment. (E) Fluorescence of HeLa cells incubated with DiI labeled exosomes from indicated cells by fluorescent microscopy. Scale bar = 50 μm. (F) Transfer purified exosomes from 293T cells co-transfected with Flag-circALTO1/2 to HeLa cell were detected by immunoblotting with the indicated antibodies. (G) Transfer purified exosomes from 293T cells co-transfected with pCDNA, Flag-circALTO1/2 to HeLa cell. After 24 hours, transfected pCDNA-GFP to the cells for 48 hours, then harvest the cells. GFP and FLAG were detected by immunoblotting. HSP90 is the loading control. (H) HeLa cells were show as in (G). Cells in PBS were imaged. Scale bar = 200 μm.

These purified EVs were used to determine whether circALTO RNAs could be transferred to HeLa cells through exosomes. EVs were purified, fluorescently labeled with DiI, and incubated with HeLa cells for 24 h. Fluorescence was only detected in HeLa cells incubated with labeled vector control or circALTO1/2 EVs, but not with PBS containing dye (Figure 6E). Next, the expression of Flag-ALTO in cell lysates was determined by immunoblotting. Flag-ALTO expression was detected in HeLa recipient cells for both circALTO vectors (Figure 6F). The expression of Flag-ALTO from circALTO1 was particularly strong, perhaps due to its potential for rolling-circle translation. Next, we tested whether circALTO RNA containing exosomes could mediate transcriptional activation as has been shown for co-transfected circALTO plasmid. Thus, exosomes were purified from pCDNA3.1, pcircALTO1, or pcircALTO2 transfected cells, and incubated with HeLa cells. Those exosome treated cells were then transfected with pCDNA-GFP and the expression of GFP was monitored by WB (Figure 6G) and IF (Figure 6H). GFP expression was notably higher in the circALTO exosome-treated cells. These data strengthen the conclusion that MCPyV ALTO promotes the expression of specific co-transfected plasmids.

### CircALTO formation is conserved in TSPyV

Like MCPyV, TSPyV is an alphapolyomavirus whose genome has the potential to encode for ALTO (13,14,56). The MT/ALTO genes appear to be conserved in a subset of *Alphapolyomaviruses* and the TSPyV early region contains canonical splice sites that could also be susceptible to backsplicing events. To determine whether TSPyV might also generate a circular ALTO RNA (Figure 7A), a minigene expression vector containing the ALTO-containing region of TSPyV was generated (Supplementary Figure S5A). The TSPyV circALTO construct was transfected into 293T cells, and a product consistent with a circALTO product was detected by RT-PCR (Figure 7B). Like MCPyV circALTO, TSPyV circALTO was completely resistant to RNase R (Supplementary Figure S5B). Using antibodies specific for TSPyV ALTO (pAbMT) (13), TSPyV ALTO protein could be detected in TSPyV circALTO transfected cells (Figure 7C). Thus, like MCPyV circALTOs, TSPyV circALTO is translated to ALTO in vitro. To determine whether TSPyV circALTO could be detected during TSPyV infections, we analyzed FFPE skin biopsies from five patients diagnosed with trichodysplasia spinulosa (TS). Biopsy samples were identified from patients with clinical and histological features consistent with TS (Figure 7D). All patients developed TS in the setting of immunosuppression (Supplementary Figure S5C). Endpoint RT-PCR revealed that products consistent with linear early region transcripts (Linear small T) could be identified in 4 out of 5 samples, and circALTO could be detected in 3 out of 5 samples (Figure 7E). Sanger sequencing confirmed that the backsplice junction in patient tissues was identical to the product generated by transfection of the TSPyV circALTO construct (Figure 7F). Finally, we tested whether TSPyV circALTO might also promote co-transfected genes. In contrast to MCPyV circALTO expression, TSPyV circALTO did not promote GFP reporter expression from the CMV as assessed by WB, IF and qRT-PCR (Supplementary Figure S5D-F). We conclude that circALTO formation is conserved in TSPyV; however, the function of TSPyV ALTO remains unknown.

**Figure 7.**
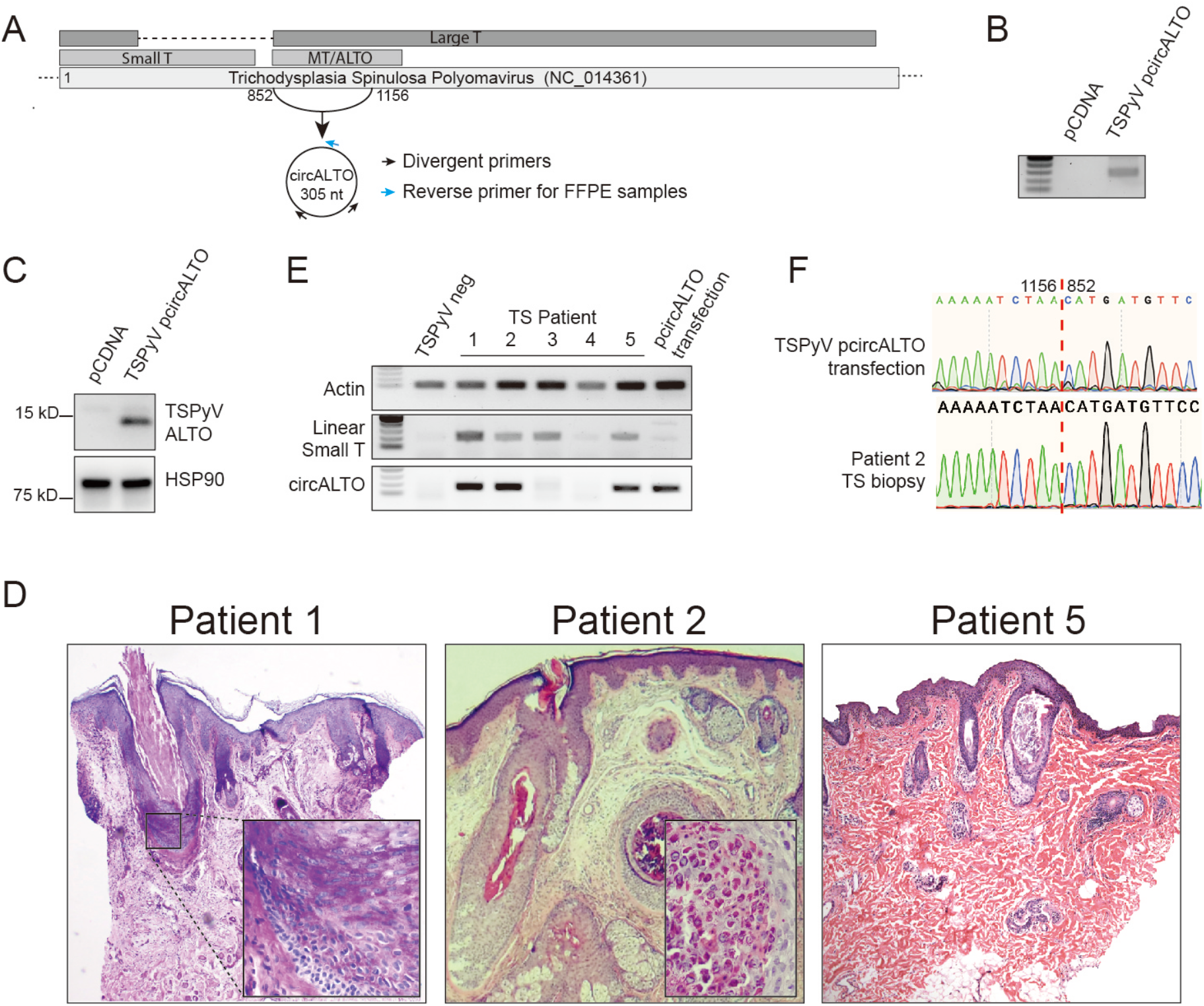
Conservation of circALTO in TSPyV. (A) Schematic representation of a predicted circALTO in TSPyV. Diagram highlights the location of the divergent primers used to detect TSPyV circALTO. (B). RT-PCR analysis of cDNA prepared from total RNA prepared with random hexamer primers. 293T cells were transfected with pCDNA vector or a construct containing the potentially backspliced early region of TSPyV. Total RNA was first treated with RNase R. (C) Western blot for TSPyV ALTO from 293T cells transfected with a pCDNA vector or the TSPyV circALTO construct. Lysates were probed with a polyclonal antibody generated against 2 peptides specific to the TSPyV ALTO, pAbMT (13). HSP90, loading control. (D) Images from cases of TS identified in this study. Clinical images (top) from TS patients 1, 2, and 5. The photos show the characteristic folliculocentric papules on the face and neck. Histological images (bottom) show the H&E stained image of lesional skin from patients 1, 2, and 5. Affected skin shows dysmorphic hair follicles. Patient 1 inset shows expansion of inner root sheath (IRS). Patient 2 inset shows enlarged trichohyaline granules in IRS cells. (E) Total RNA was extracted from lesional biopies and cDNA prepared with random hexamer primers. RT-PCR analysis using indicated primers revealed the presence of linear small T (sT) in 4/5 TS samples and circALTO in 3/5 TS samples. A TSPyV negative squamous cell carcinoma was used as a negative control. 293T cells transfected with TSPyV circALTO was used as a positive control for circALTO. (F) Sanger sequencing of PCR products from 293T transfected cells and PCR product from patient 2 confirmed the expected backsplice junction.

## DISCUSSION

EBV, KSHV, and HPV have been shown to encode for virus-derived circRNAs (24,25,27-29). In this report, we reveal the existence of polyomavirus-encoded circRNAs. We characterize both MCPyV and TSPyV-derived, protein-coding circALTO RNAs. MCPyV circALTOs can be detected in both VP-MCC lines and patient tumors. RNase R digestion and northern blotting confirm that the products are bona fide circRNAs, as opposed to RNA transcripts making multiple passes around the circular viral genome. The biological relevance of circALTO is strengthened by our identification and detection of TSPyV circALTO in TS patient samples, suggesting the evolutionary conservation of circALTO formation. While we focus only on circALTO, our analyses suggest that backsplicing in HPyV is much more widespread than previously appreciated and may involve other genes, such as those encoding agnoproteins.

We provide conclusive evidence that circALTOs function as templates for translation. Small interfering RNAs specifically targeting the unique backsplice (circularization) junction, but not siRNAs specific to the linear ALTO RNAs, dramatically decrease ALTO protein expression in vitro. Moreover, mutation of in the MCPyV circALTO backsplicing sites almost completely abrogates expression of the ALTO protein isoform.

The protein coding capacity of TSPyV circALTO is also conserved and was confirmed using MT/ALTO specific antibodies in vitro. Both MCPyV and TSPyV also generate a diverse array of linear transcripts that also have the potential to encode for ALTO. Though linear and circular RNAs possess different properties, it remains unclear how these multiple transcripts are differentially regulated and whether they contribute uniquely to viral replication.

The factors that regulate the cap-independent translation of circRNAs remain an area of active investigation. Internal ribosome entry sequences, m^6^A motifs, and novel sequences in the untranslated region (UTR) of circRNAs have all been demonstrated to promote circRNA translation (57,58). While circALTOs are m^6^A modified, the knockdown of Mettl3 and subsequent decrease of m^6^A levels do not obviously impact ALTO protein levels (Supplementary Figure S3D-G). Thus, we speculate that m^6^A modification might facilitate the biogenesis, rather than translation, of circALTOs. As m^6^A has been reported to play diverse roles in splicing (45), translation (44), and circularization of circRNAs (58), it likely that the function of this abundant modification may be multifactorial and context specific.

CircALTO RNAs are perfectly complementary to the MCPyV encoded miRNA, miR-M1-5p/3p. However, because circALTO is much less abundant relative to linear ALTO mRNAs, it is unlikely that a primary function of circALTO would be to function as a decoy or sponge for miRNAs, as has been reported for other circRNAs (10,22,30,31). Indeed, our experiments did not suggest that circALTO could function as a sponge for miRNAs. Rather, we find that MCPyV miRNAs actually have the potential to regulate circALTO expression in co-transfection experiments. While there is clear biochemical evidence that a subset of circRNAs sponge and inhibit specific miRNAs (30,31,48), we contend that the low abundance of most circRNAs makes it unlikely that miRNA sponging is a biologically relevant function for the majority of circRNAs. Certainly, the complex dynamics of the relationship between circALTOs and miRNAs during polyomavirus infections requires additional investigation.

MPyV MT enhances the transcription genes whose promoters contain binding sites for the polyomavirus enhancer A binding proteins 1 and 3 (PEA1/AP-1) and (PEA3/ETV4). These promoters are also required for MPyV replication (59,60). In addition, SV40 large T antigen has previously been reported to promote transcription from some viral and cellular promoters (61,62). For the first time we demonstrate a biological function for MCPyV ALTO and show that it can, like MPyV MT, amplify the transcriptional activity of some promoters. Specifically, circALTO that encodes for ALTO, but not a splicing mutant or start-codon mutant, dramatically enhanced the transcription of reporter genes driven by the viral CMV promoter, but not the human EF1-*α* promoter, in co-transfected plasmids. While SV40 LT has been shown to bind multiple components of the transcriptional pre-initiation complex, the precise mechanism of its ability to promote transcription remains unclear (63). Additional studies will be necessary to identify the promoter sequence elements and potential interacting proteins that might regulate ALTO’s ability to promote the activity of specific promoters. Curiously, while circALTO formation appears to be conserved, TSPyV ALTO’s ability to promote transcription, at least in the two promoters tested here, did not appear to be conserved. While the upregulation of viral promoters suggests an obvious avenue through which ALTO might promote viral replication, additional studies are necessary to determine whether ALTO plays an essential role in the replication of MCPyV or TSPyV in vitro or in vivo.

Finally, we provide strong evidence that circALTO RNAs can be enriched in exosomes. While the exosome-mediated transfer and translation of linear mRNAs between cells has previously been documented, we present the first evidence of circRNA transfer between cells through EVs with exosomal characteristics. Consistent with the literature (35), we found circALTO to be enriched in exosomes relative to linear ALTO compared to their levels in the cell. Coupled with our confirmation that circALTO is far more stable than linear ALTO, we speculate that exosomal circALTO could be transferred to recipient cells and promote transcription in a paracrine fashion. A speculative hypothesis might be that circALTO RNA-containing exosomes could promote MCPyV infection in surrounding tissues by “ priming” neighboring to reduce the immune response and/or promote MCPyV gene transcription and replication (Supplementary Figure S6).

Exosomes also have diagnostic potential in cancer (36). We were able to detect circALTO in WaGa cells. EVs, including exosomes, have been found in serum and variety of other bodily fluids. Thus, despite the low levels of circALTO in WaGa cells, it would nonetheless be worthwhile determining whether circALTO might have utility as a non-invasive clinical biomarker for patient with VP-MCC.

Our discovery of circALTO offer novel insights on the transcriptional complexity of polyomaviruses and extends the spectrum of viruses that encode for circRNAs. The incorporation of circRNAs into EVs, which can be transferred and expressed in other cells, raises the possibility that circALTO exerts its effects primarily in a non-cell-autonomous fashion. While MCPyV ALTO can regulate transcription, the precise role of MCPyV and TSPyV circALTO in models of infection will require further investigation. Additional studies on ALTO may yield novel therapeutic targets in HPyV infection or carcinogenesis and will broaden our understanding of the regulation of circRNA metabolism and the polyomavirus life cycle.

## ACKNOWLEDGEMENTS

Smaranda Willcox and Dr. Jack Griffith (University of North Carolina) are thanked for performing the EM. Dr. Christine Ko (Yale) is thanked for providing TS patient tissues and histologic images. Dr. Christopher Buck (NIH) is thanked for providing pU-5864 and comments on the manuscript.

## CONTRIBUTIONS

R.W. and R.Y. conceived and designed the experiments; R.Y. performed most of the experiments with assistance from E.L., E.K., and R.W.; J.K. analyzed RNA-seq data; C.C. performed NTA assays; J.C., Y.T., and C.S. designed, performed, and analyzed miRNA and luciferase assays. C.C., T.S., L.R., and M.F. provided critical reagents. R.Y. and R.W. wrote the manuscript with input from all co-authors.

## FUNDING

This work was supported by grants from the UT Southwestern Cary Council, ACS (RSG-18-058-01), and NIAMS (R01AR072655) to R.W.; and a Burroughs Wellcome Investigators in Pathogenesis and Infectious Disease Award (#1011070) and NIAID (R21AI147178) to C.S.S.

## DATA AVAILABILITY

All data supporting the findings of this study are available from the corresponding author upon reasonable request.

## Notes

### Competing Interest Statement

The authors have declared no competing interest.

## References

1. Nguyen, K.D., Chamseddin, B.H., Cockerell, C.J. and Wang, R.C. (2019) The Biology and Clinical Features of Cutaneous Polyomaviruses. J Invest Dermatol, 139, 285–292.

2. Feng, H., Shuda, M., Chang, Y. and Moore, P.S. (2008) Clonal integration of a polyomavirus in human Merkel cell carcinoma. Science, 319, 1096–1100.

3. van der Meijden, E., Janssens, R.W., Lauber, C., Bouwes Bavinck, J.N., Gorbalenya, A.E. and Feltkamp, M.C. (2010) Discovery of a new human polyomavirus associated with trichodysplasia spinulosa in an immunocompromized patient. PLoS Pathog, 6, e1001024.

4. Matthews, M.R., Wang, R.C., Reddick, R.L., Saldivar, V.A. and Browning, J.C. (2011) Viral-associated trichodysplasia spinulosa: a case with electron microscopic and molecular detection of the trichodysplasia spinulosa-associated human polyomavirus. Journal of Cutaneous Pathology, 38, 420–431.

5. Kazem, S., van der Meijden, E., Wang, R.C., Rosenberg, A.S., Pope, E., Benoit, T., Fleckman, P. and Feltkamp, M.C. (2014) Polyomavirus-associated Trichodysplasia spinulosa involves hyperproliferation, pRB phosphorylation and upregulation of p16 and p21. PLoS One, 9, e108947.

6. DeCaprio, J.A. and Garcea, R.L. (2013) A cornucopia of human polyomaviruses. Nature reviews. Microbiology, 11, 264–276.

7. Houben, R., Shuda, M., Weinkam, R., Schrama, D., Feng, H., Chang, Y., Moore, P.S. and Becker, J.C. (2010) Merkel cell polyomavirus-infected Merkel cell carcinoma cells require expression of viral T antigens. Journal of virology, 84, 7064–7072.

8. Moens, U., Krumbholz, A., Ehlers, B., Zell, R., Johne, R., Calvignac-Spencer, S. and Lauber, C. (2017) Biology, evolution, and medical importance of polyomaviruses: An update. Infect Genet Evol, 54, 18–38.

9. Schowalter, R.M. and Buck, C.B. (2013) The Merkel cell polyomavirus minor capsid protein. PLoS Pathog, 9, e1003558.

10. Seo, G.J., Chen, C.J. and Sullivan, C.S. (2009) Merkel cell polyomavirus encodes a microRNA with the ability to autoregulate viral gene expression. Virology, 383, 183–187.

11. Lee, S., Paulson, K.G., Murchison, E.P., Afanasiev, O.K., Alkan, C., Leonard, J.H., Byrd, D.R., Hannon, G.J. and Nghiem, P. (2011) Identification and validation of a novel mature microRNA encoded by the Merkel cell polyomavirus in human Merkel cell carcinomas. Journal of clinical virology: the official publication of the Pan American Society for Clinical Virology, 52, 272–275.

12. Carter, J.J., Daugherty, M.D., Qi, X., Bheda-Malge, A., Wipf, G.C., Robinson, K., Roman, A., Malik, H.S. and Galloway, D.A. (2013) Identification of an overprinting gene in Merkel cell polyomavirus provides evolutionary insight into the birth of viral genes. Proc Natl Acad Sci U S A, 110, 12744–12749.

13. van der Meijden, E., Kazem, S., Dargel, C.A., van Vuren, N., Hensbergen, P.J. and Feltkamp, M.C. (2015) Characterization of T Antigens, Including Middle T and Alternative T, Expressed by the Human Polyomavirus Associated with Trichodysplasia Spinulosa. Journal of virology, 89, 9427–9439.

14. van der Meijden, E. and Feltkamp, M. (2018) The Human Polyomavirus Middle and Alternative T-Antigens; Thoughts on Roles and Relevance to Cancer. Front Microbiol, 9, 398.

15. Cheng, J., DeCaprio, J.A., Fluck, M.M. and Schaffhausen, B.S. (2009) Cellular transformation by Simian Virus 40 and Murine Polyoma Virus T antigens. Seminars in cancer biology, 19, 218–228.

16. Kristensen, L.S., Andersen, M.S., Stagsted, L.V.W., Ebbesen, K.K., Hansen, T.B. and Kjems, J. (2019) The biogenesis, biology and characterization of circular RNAs. Nat Rev Genet, 20, 675–691.

17. Li, X., Yang, L. and Chen, L.L. (2018) The Biogenesis, Functions, and Challenges of Circular RNAs. Mol Cell, 71, 428–442.

18. Sanger, H.L., Klotz, G., Riesner, D., Gross, H.J. and Kleinschmidt, A.K. (1976) Viroids are single-stranded covalently closed circular RNA molecules existing as highly base-paired rod-like structures. Proc Natl Acad Sci U S A, 73, 3852–3856.

19. Hsu, M.T. and Coca-Prados, M. (1979) Electron microscopic evidence for the circular form of RNA in the cytoplasm of eukaryotic cells. Nature, 280, 339–340.

20. Conn, V.M., Hugouvieux, V., Nayak, A., Conos, S.A., Capovilla, G., Cildir, G., Jourdain, A., Tergaonkar, V., Schmid, M., Zubieta, C. et al. (2017) A circRNA from SEPALLATA3 regulates splicing of its cognate mRNA through R-loop formation. Nat Plants, 3, 17053.

21. Danan, M., Schwartz, S., Edelheit, S. and Sorek, R. (2012) Transcriptome-wide discovery of circular RNAs in Archaea. Nucleic Acids Res, 40, 3131–3142.

22. Han, D., Li, J., Wang, H., Su, X., Hou, J., Gu, Y., Qian, C., Lin, Y., Liu, X., Huang, M. et al. (2017) Circular RNA circMTO1 acts as the sponge of microRNA-9 to suppress hepatocellular carcinoma progression. Hepatology, 66, 1151–1164.

23. Yu, J., Xu, Q.G., Wang, Z.G., Yang, Y., Zhang, L., Ma, J.Z., Sun, S.H., Yang, F. and Zhou, W.P. (2018) Circular RNA cSMARCA5 inhibits growth and metastasis in hepatocellular carcinoma. J Hepatol, 68, 1214–1227.

24. Toptan, T., Abere, B., Nalesnik, M.A., Swerdlow, S.H., Ranganathan, S., Lee, N., Shair, K.H., Moore, P.S. and Chang, Y. (2018) Circular DNA tumor viruses make circular RNAs. Proc Natl Acad Sci U S A.

25. Ungerleider, N., Concha, M., Lin, Z., Roberts, C., Wang, X., Cao, S., Baddoo, M., Moss, W.N., Yu, Y., Seddon, M. et al. (2018) The Epstein Barr virus circRNAome. PLoS Pathog, 14, e1007206.

26. Chamseddin, B.H., Lee, E.E., Kim, J., Zhan, X., Yang, R., Murphy, K.M., Lewis, C., Hosler, G.A., Hammer, S.T. and Wang, R.C. (2019) Assessment of circularized E7 RNA, GLUT1, and PD-L1 in anal squamous cell carcinoma. Oncotarget, 10, 5958–5969.

27. Abere, B., Li, J., Zhou, H., Toptan, T., Moore, P.S. and Chang, Y. (2020) Kaposi’s Sarcoma-Associated Herpesvirus-Encoded circRNAs Are Expressed in Infected Tumor Tissues and Are Incorporated into Virions. mBio, 11.

28. Tagawa, T., Gao, S., Koparde, V.N., Gonzalez, M., Spouge, J.L., Serquina, A.P., Lurain, K., Ramaswami, R., Uldrick, T.S., Yarchoan, R. et al. (2018) Discovery of Kaposi’s sarcoma herpesvirus-encoded circular RNAs and a human antiviral circular RNA. Proc Natl Acad Sci U S A, 115, 12805–12810.

29. Zhao, J., Lee, E.E., Kim, J., Yang, R., Chamseddin, B., Ni, C., Gusho, E., Xie, Y., Chiang, C.M., Buszczak, M. et al. (2019) Transforming activity of an oncoprotein-encoding circular RNA from human papillomavirus. Nature communications, 10, 2300.

30. Zheng, Q., Bao, C., Guo, W., Li, S., Chen, J., Chen, B., Luo, Y., Lyu, D., Li, Y., Shi, G. et al. (2016) Circular RNA profiling reveals an abundant circHIPK3 that regulates cell growth by sponging multiple miRNAs. Nature communications, 7, 11215.

31. Hansen, T.B., Jensen, T.I., Clausen, B.H., Bramsen, J.B., Finsen, B., Damgaard, C.K. and Kjems, J. (2013) Natural RNA circles function as efficient microRNA sponges. Nature, 495, 384–388.

32. Abdelmohsen, K., Panda, A.C., Munk, R., Grammatikakis, I., Dudekula, D.B., De, S., Kim, J., Noh, J.H., Kim, K.M., Martindale, J.L. et al. (2017) Identification of HuR target circular RNAs uncovers suppression of PABPN1 translation by CircPABPN1. RNA Biol, 14, 361–369.

33. Zhang, Y., Zhang, X.O., Chen, T., Xiang, J.F., Yin, Q.F., Xing, Y.H., Zhu, S., Yang, L. and Chen, L.L. (2013) Circular intronic long noncoding RNAs. Mol Cell, 51, 792–806.

34. Legnini, I., Di Timoteo, G., Rossi, F., Morlando, M., Briganti, F., Sthandier, O., Fatica, A., Santini, T., Andronache, A., Wade, M. et al. (2017) Circ-ZNF609 Is a Circular RNA that Can Be Translated and Functions in Myogenesis. Mol Cell, 66, 22–37 e29.

35. Li, Y., Zheng, Q., Bao, C., Li, S., Guo, W., Zhao, J., Chen, D., Gu, J., He, X. and Huang, S. (2015) Circular RNA is enriched and stable in exosomes: a promising biomarker for cancer diagnosis. Cell Res, 25, 981–984.

36. Wang, Y., Liu, J., Ma, J., Sun, T., Zhou, Q., Wang, W., Wang, G., Wu, P., Wang, H., Jiang, L. et al. (2019) Exosomal circRNAs: biogenesis, effect and application in human diseases. Mol Cancer, 18, 116.

37. Theiss, J.M., Gunther, T., Alawi, M., Neumann, F., Tessmer, U., Fischer, N. and Grundhoff, A. (2015) A Comprehensive Analysis of Replicating Merkel Cell Polyomavirus Genomes Delineates the Viral Transcription Program and Suggests a Role for mcv-miR-M1 in Episomal Persistence. PLoS Pathog, 11, e1004974.

38. Daily, K., Coxon, A., Williams, J.S., Lee, C.R., Coit, D.G., Busam, K.J. and Brownell, I. (2015) Assessment of cancer cell line representativeness using microarrays for Merkel cell carcinoma. J Invest Dermatol, 135, 1138–1146.

39. Zhao, J., Jia, Y., Shen, S., Kim, J., Wang, X., Lee, E., Brownell, I., Cho-Vega, J.H., Lewis, C., Homsi, J. et al. (2020) Merkel Cell Polyomavirus Small T Antigen Activates Noncanonical NF-kappaB Signaling to Promote Tumorigenesis. Mol Cancer Res, 18, 1623–1637.

40. Buck, C.B., Pastrana, D.V., Lowy, D.R. and Schiller, J.T. (2004) Efficient intracellular assembly of papillomaviral vectors. Journal of virology, 78, 751–757.

41. Pendleton, K.E., Chen, B., Liu, K., Hunter, O.V., Xie, Y., Tu, B.P. and Conrad, N.K. (2017) The U6 snRNA m(6)A Methyltransferase METTL16 Regulates SAM Synthetase Intron Retention. Cell, 169, 824–835 e814.

42. Jeppesen, D.K., Fenix, A.M., Franklin, J.L., Higginbotham, J.N., Zhang, Q., Zimmerman, L.J., Liebler, D.C., Ping, J., Liu, Q., Evans, R. et al. (2019) Reassessment of Exosome Composition. Cell, 177, 428–445 e418.

43. Meckes, D.G., Jr., Shair, K.H., Marquitz, A.R., Kung, C.P., Edwards, R.H. and Raab-Traub, N. (2010) Human tumor virus utilizes exosomes for intercellular communication. Proc Natl Acad Sci U S A, 107, 20370–20375.

44. Yang, Y., Fan, X., Mao, M., Song, X., Wu, P., Zhang, Y., Jin, Y., Yang, Y., Chen, L.L., Wang, Y. et al. (2017) Extensive translation of circular RNAs driven by N(6)-methyladenosine. Cell Res, 27, 626–641.

45. Price, A.M., Hayer, K.E., McIntyre, A.B.R., Gokhale, N.S., Abebe, J.S., Della Fera, A.N., Mason, C.E., Horner, S.M., Wilson, A.C., Depledge, D.P. et al. (2020) Direct RNA sequencing reveals m(6)A modifications on adenovirus RNA are necessary for efficient splicing. Nature communications, 11, 6016.

46. Zhou, C., Molinie, B., Daneshvar, K., Pondick, J.V., Wang, J., Van Wittenberghe, N., Xing, Y., Giallourakis, C.C. and Mullen, A.C. (2017) Genome-Wide Maps of m6A circRNAs Identify Widespread and Cell-Type-Specific Methylation Patterns that Are Distinct from mRNAs. Cell Rep, 20, 2262–2276.

47. Conn, S.J., Pillman, K.A., Toubia, J., Conn, V.M., Salmanidis, M., Phillips, C.A., Roslan, S., Schreiber, A.W., Gregory, P.A. and Goodall, G.J. (2015) The RNA binding protein quaking regulates formation of circRNAs. Cell, 160, 1125–1134.

48. Memczak, S., Jens, M., Elefsinioti, A., Torti, F., Krueger, J., Rybak, A., Maier, L., Mackowiak, S.D., Gregersen, L.H., Munschauer, M. et al. (2013) Circular RNAs are a large class of animal RNAs with regulatory potency. Nature, 495, 333–338.

49. Piwecka, M., Glazar, P., Hernandez-Miranda, L.R., Memczak, S., Wolf, S.A., Rybak-Wolf, A., Filipchyk, A., Klironomos, F., Cerda Jara, C.A., Fenske, P. et al. (2017) Loss of a mammalian circular RNA locus causes miRNA deregulation and affects brain function. Science, 357.

50. Liu, J., Carmell, M.A., Rivas, F.V., Marsden, C.G., Thomson, J.M., Song, J.J., Hammond, S.M., Joshua-Tor, L. and Hannon, G.J. (2004) Argonaute2 is the catalytic engine of mammalian RNAi. Science, 305, 1437–1441.

51. O’Hara, B.A., Morris-Love, J., Gee, G.V., Haley, S.A. and Atwood, W.J. (2020) JC Virus infected choroid plexus epithelial cells produce extracellular vesicles that infect glial cells independently of the virus attachment receptor. PLoS Pathog, 16, e1008371.

52. Morris-Love, J., Gee, G.V., O’Hara, B.A., Assetta, B., Atkinson, A.L., Dugan, A.S., Haley, S.A. and Atwood, W.J. (2019) JC Polyomavirus Uses Extracellular Vesicles To Infect Target Cells. mBio, 10.

53. Crewe, C., Joffin, N., Rutkowski, J.M., Kim, M., Zhang, F., Towler, D.A., Gordillo, R. and Scherer, P.E. (2018) An Endothelial-to-Adipocyte Extracellular Vesicle Axis Governed by Metabolic State. Cell, 175, 695–708 e613.

54. Pegtel, D.M., Cosmopoulos, K., Thorley-Lawson, D.A., van Eijndhoven, M.A., Hopmans, E.S., Lindenberg, J.L., de Gruijl, T.D., Wurdinger, T. and Middeldorp, J.M. (2010) Functional delivery of viral miRNAs via exosomes. Proc Natl Acad Sci U S A, 107, 6328–6333.

55. Valadi, H., Ekstrom, K., Bossios, A., Sjostrand, M., Lee, J.J. and Lotvall, J.O. (2007) Exosome-mediated transfer of mRNAs and microRNAs is a novel mechanism of genetic exchange between cells. Nat Cell Biol, 9, 654–659.

56. Kazem, S., Lauber, C., van der Meijden, E., Kooijman, S., Kravchenko, A.A., TrichSpin, N., Feltkamp, M.C. and Gorbalenya, A.E. (2016) Limited variation during circulation of a polyomavirus in the human population involves the COCO-VA toggling site of Middle and Alternative T-antigen(s). Virology, 487, 129–140.

57. Pamudurti, N.R., Bartok, O., Jens, M., Ashwal-Fluss, R., Stottmeister, C., Ruhe, L., Hanan, M., Wyler, E., Perez-Hernandez, D., Ramberger, E. et al. (2017) Translation of CircRNAs. Mol Cell, 66, 9–21 e27.

58. Tang, C., Xie, Y., Yu, T., Liu, N., Wang, Z., Woolsey, R.J., Tang, Y., Zhang, X., Qin, W., Zhang, Y. et al. (2020) m(6)A-dependent biogenesis of circular RNAs in male germ cells. Cell Res, 30, 211–228.

59. Gottlieb, K.A. and Villarreal, L.P. (2001) Natural biology of polyomavirus middle T antigen. Microbiol Mol Biol Rev, 65, 288-318; second and third pages, table of contents.

60. Wasylyk, C., Imler, J.L. and Wasylyk, B. (1988) Transforming but not immortalizing oncogenes activate the transcription factor PEA1. EMBO J, 7, 2475–2483.

61. Rice, P.W. and Cole, C.N. (1993) Efficient transcriptional activation of many simple modular promoters by simian virus 40 large T antigen. Journal of virology, 67, 6689–6697.

62. Gilinger, G. and Alwine, J.C. (1993) Transcriptional activation by simian virus 40 large T antigen: requirements for simple promoter structures containing either TATA or initiator elements with variable upstream factor binding sites. Journal of virology, 67, 6682–6688.

63. Johnston, S.D., Yu, X.M. and Mertz, J.E. (1996) The major transcriptional transactivation domain of simian virus 40 large T antigen associates nonconcurrently with multiple components of the transcriptional preinitiation complex. Journal of virology, 70, 1191–1202.

