## Supplementary Figures for "Human Polyomavirus-Encoded Circular RNAs"

A

| Sample | Chromosome | Breakpoint 1 | Breakpoint 2 | Strand | Junction read count | Junction read ratio |
|---|---|---|---|---|---|---|
| ERS760222 | NC_010277.2 | 4950 | 5113 | + | 15 | 53.6% |
| ERS760222 | NC_010277.2 | 666 | 3142 | – | 1 | 0.3% |
| ERS760222 | NC_010277.2 | 666 | 1427 | – | 2 | 0.4% |
| ERS760223 | NC_010277.2 | 4950 | 5113 | + | 27 | 36.7% |
| ERS760223 | NC_010277.2 | 666 | 3142 | – | 1 | 0.4% |
| ERS760223 | NC_010277.2 | 666 | 2760 | – | 2 | 0.8% |
| ERS760224 | NC_010277.2 | 4950 | 5113 | + | 3 | 54.5% |
| ERS760224 | NC_010277.2 | 4950 | 5035 | + | 1 | 22.2% |
| ERS760224 | NC_010277.2 | 666 | 3142 | – | 1 | 0.2% |
| ERS760224 | NC_010277.2 | 666 | 1427 | – | 1 | 0.2% |
| ERS760225 | NC_010277.2 | 4950 | 5113 | + | 5 | 37.0% |
| ERS760225 | NC_010277.2 | 666 | 1427 | – | 1 | 0.3% |

B

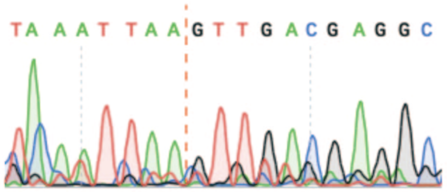


C

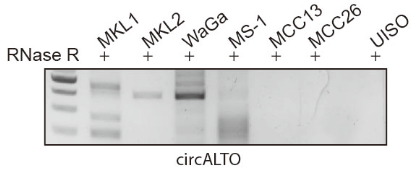


D

| Specimen Label | Anatomic Site | Pathological status | Neo Plastic Cellularity Percentage |
|---|---|---|---|
| 26974-Frozen-5 | Connective, subcutaneous and other soft tissues | Malignant | 90 |
| 24902-Frozen-6 | Lymph node, NOS | Malignant | 80 |
| 28548-Frozen-5 | Parotid gland | Malignant | 80 |
| 26579-Frozen-3 | Skin, NOS | Malignant | 60 |
| 24118-Frozen-4 | Skin, NOS | Malignant | 80 |
| 25376-Frozen-4 | Lymph node, NOS | Malignant | 50 |
| 30312-Frozen-3 | Skin, NOS | Non-Malignant | |

E

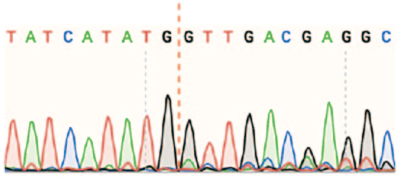


**Supplementary Figure S1. Identification of MCPyV circRNAs.**
(A) Table of MCPyV circRNAs identified in SRA datasets containing location of putative circRNA, read count, and backsplice ratio.
(B) Sanger sequencing of the non-specific band between the circALTO1 and circALTO2 products from WaGa cells show the inclusion of non-MCPyV sequence at the backsplice junction.
(C) Endpoint RT-PCR analysis of circALTOs from VP-MCC lines (MKL-1, MKL-2, MS1 and WaGa) and VN-MCC (MCC13, MCC26 and UISO). Total RNA treated with RNase R.
(D) Pathological features of the patient MCC samples.
(E) Sanger sequencing of PCR products from the sample No. 28548 confirmed the expected backsplice junction for circALTO2.

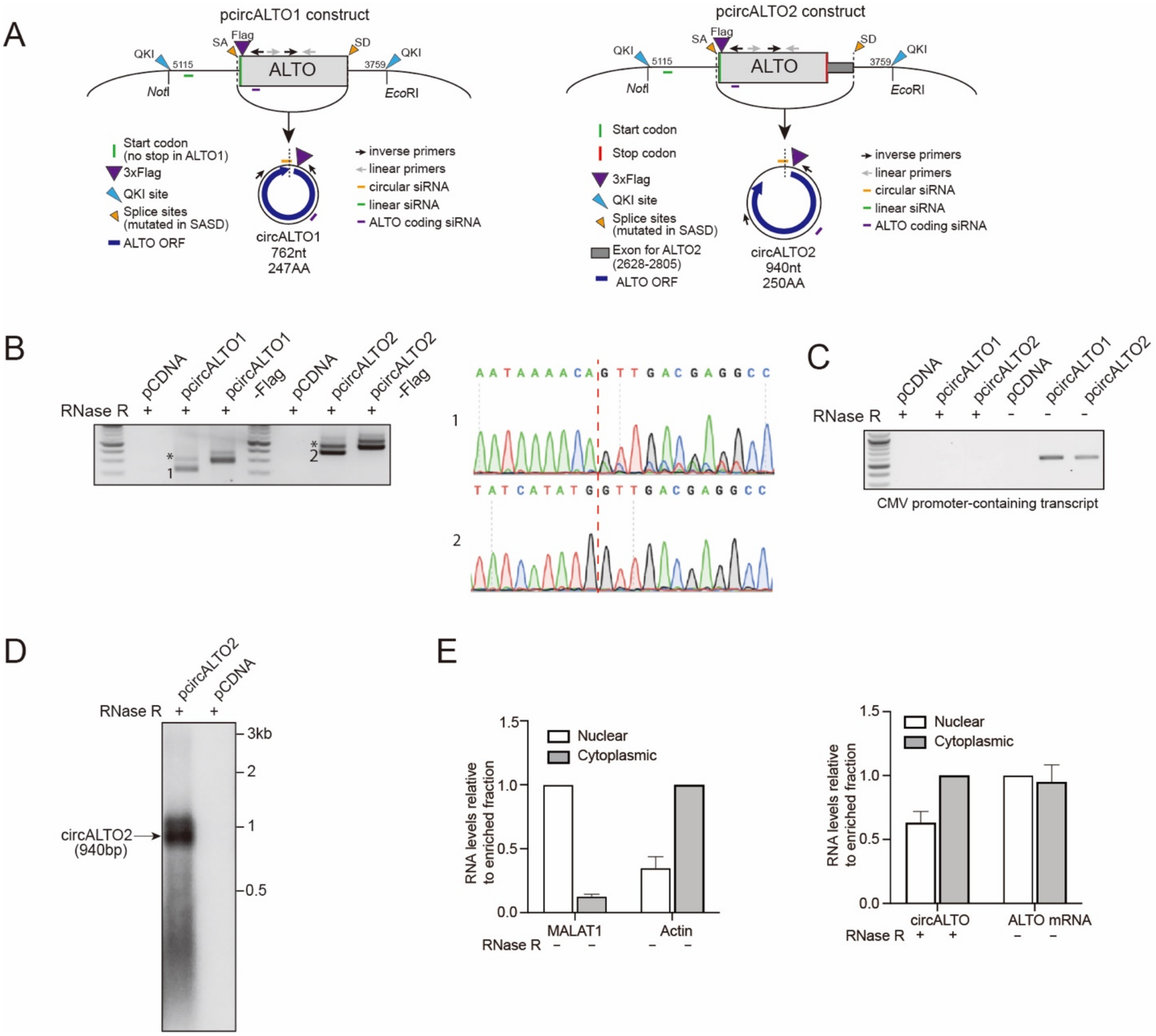
A
pcircALTO1 construct
QKI
5115
SA
Flag
ALTO
SD
3759
QKI
NotI
EcoRI
Start codon
(no stop in ALTO1)
3xFlag
QKI site
Splice sites
(mutated in SASD)
ALTO ORF
circALTO1
762nt
247AA
inverse primers
linear primers
circular siRNA
linear siRNA
ALTO coding siRNA
pcircALTO2 construct
Start codon
Stop codon
3xFlag
QKI site
Splice sites
(mutated in SASD)
Exon for ALTO2
(2628-2805)
ALTO ORF
circALTO2
940nt
250AA
B
RNase R
pCDNA
pcircALTO1
pcircALTO1-Flag
pcircALTO2
pcircALTO2-Flag
AATAAAACAGTTGACGAGGCC
TATCATATGGTTGACGAGGCC
C
CMV promoter-containing transcript
D
circALTO2
(940bp)
3kb
2
1
0.5
E
RNA levels relative to enriched fraction
Nuclear
Cytoplasmic
MALAT1
Actin
circALTO
ALTO mRNA

**Supplementary Figure S2. Chacterization of circALTO constructs.**
(A) Schematic diagram of circALTO1 (Left panel) and circALTO2 (Right panel) expression constructs generated in vitro. The location of QKI sites, ORF, 3xFLAG epitope-tag (present in ‘FLAG’), mutated splice sites (short for ‘SASD’), and siRNAs used in subsequent knockdown assay were indicated in the diagram.
(B) The formation of circALTOs from 293T cells co-transfected with pcDNA3.1-circALTO1/2 constructs were confirmed by RT-PCR (Left panel). Sanger sequencing of PCR products showed the non-specific backsplice junction with the insertion of additional nucleotides (Right panel).
(C) Readthrough transcript from 293T cells co-transfected with pcDNA3.1-circALTO1/2 constructs with or without RNase R treatment were analyzed by RT-PCR. Primers designed from CMV promoter region to the regions that flanked circALTOs.
(D) Northern blot of total RNA from 293T cells co-transfected with vector control and pcDNA3.1-circALTO2 constructs probed with linear ALTO after RNase R treatment. Arrows indicates RNase R resistant band circALTO2.
(E) Nuclear and cytoplasmic fractionation assay of 293 T cells co-transfected with circALTO1 construct performed by qRT-PCR. MALAT1 and *ACTB* served as fractionation controls. Values normalized to the enriched fraction. Error bars represent the SD of three biological replicates.

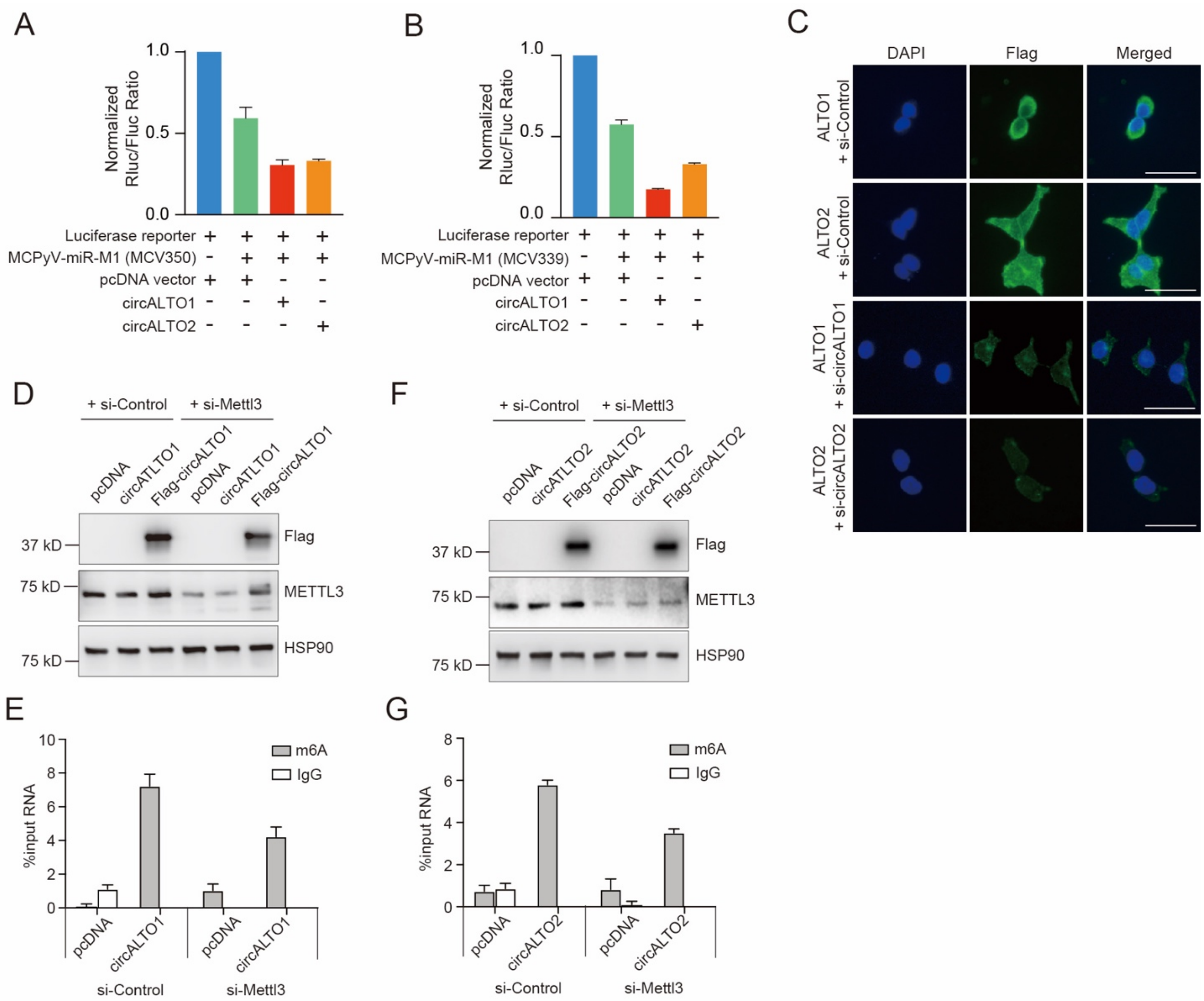


**Supplementary Figure S3. Characterization of circALTOs with siRNAs.**

(A) 293T cells were transfected with Renilla luciferase reporter with MCPyV miRNA (MCV350) and the indicated plasmids including pcDNA3.1 control vector, pcDNA circALTO1 or pcDNA circALTO2 expression plasmid. Firefly luciferase served as a transfection control and Renilla luciferase levels are plotted normalized relative to firefly luciferase levels (n=2 biological replicates).

(B) 293 cells were transfected with Renilla luciferase reporter with MCPyV miRNA (MCV339) and the indicated plasmids. n=2 biological replicates.

(C) 293T cells were transfected with the Flag-circALTO1/2 plasmids alone and Flag-circALTO1/2 with indicated siRNAs. After 48 h of transfection, the cells were fixed and stained for FLAG (green), and DAPI (blue). Scale bar = 50 μm.

(D) Western blots for METTL3 and Flag from 293T co-transfected with control or METTL3 siRNA and circALTO1 construct. HSP90, loading control.

(E) qRT-PCR of m6A or IgG control RIP of 293T co-transfected with control or METTL3 siRNA and circALTO1 construct. SON, m6A RNA IP control. Error bars represent the SD (n = 3).

(F) Western blots for METTL3 and Flag from 293T co-transfected with indicated siRNA and circALTO2. HSP90, loading control.

(G) qRT-PCR of m6A or IgG control RIP of 293T co-transfected with specific siRNA and circALTO2. SON, m6A RNA IP control. Error bars represent the SD (n = 3).

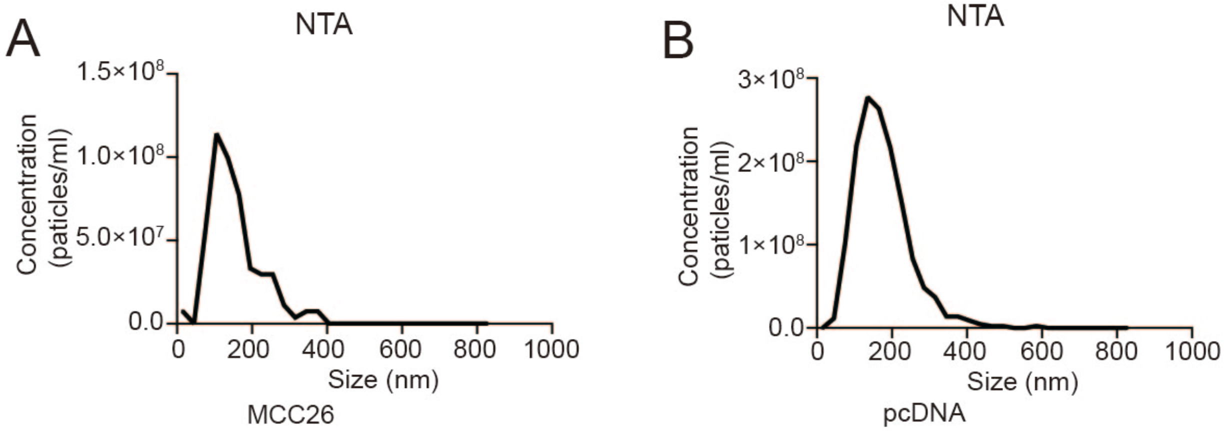


**Supplementary Figure S4. Characterization of exosomes from MCC26 and 293T cells co-transfected with pcDNA vector.**

(A-B) Size distribution analysis of total exosomes isolated from MCC26 cells (A) and 293T cells co-transfected with pcDNA vector (B).

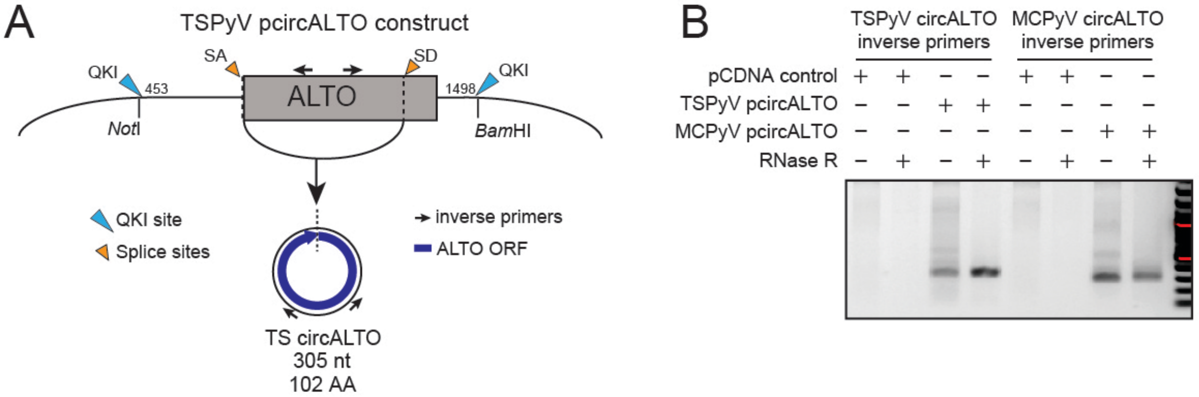


C

| Case | Sex | Age | Medical History | Onset | Linear ST | circALTO | Reference |
|---|---|---|---|---|---|---|---|
| **1** | F | 43 | Kidney transplant on tacrolimus, mycophenolate, and prednisone | 2012 | + | + | Kazem et al. (6) |
| **2** | F | 41 | Kidney transplant on mycophenolate, methylprednisolone, tacrolimus | 2017 | + | + | This study |
| **3** | F | 30 | Kidney transplant | 2019 | + | -/+ | This study |
| **4** | F | 51 | ALL | 2016 | -/+ | - | This study |
| **5** | M | 7 | Pre-B cell ALL treated with methotrexate, 6-MP, vincristine | 2018 | + | + | This study |

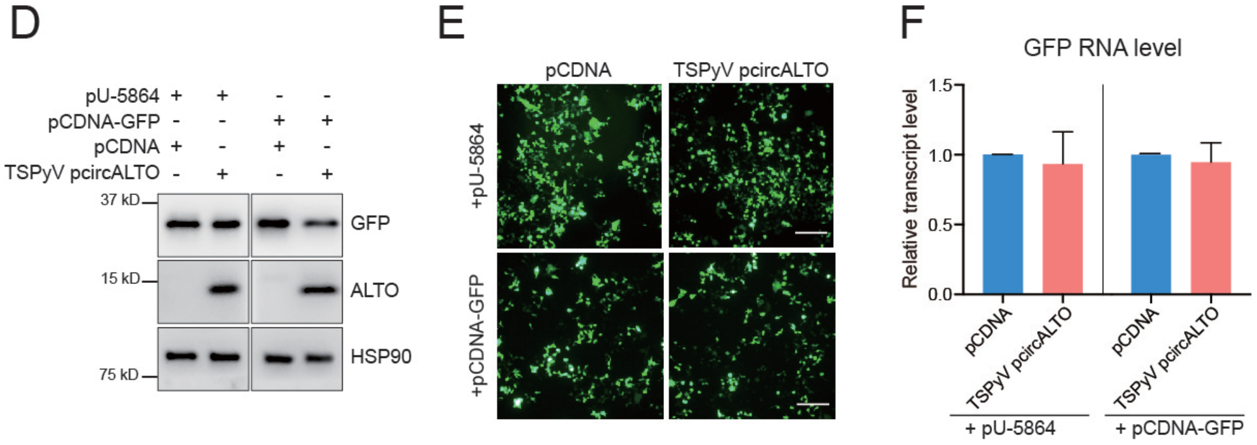

**Supplementary Figure S5. Identification and characterization of TSPyV circALTO.**
(A) Schematic diagram of TSPyV circALTO expression constructs generated in vitro. The location of QKI sites, ORF, and splice sites were indicated in the diagram.
(B) RT-PCR analysis of total RNA from 293T cells co-transfected with MCPyV circALTO or TSPyV circALTO plasmids with and without RNase R treatment.
(C) Table summarizing the clinical characteristics and RT-PCR results of TS patients utilized in this study. Only patient 1 has previously been reported.
(D) Western blot for GFP and ALTO from 293T cells were co-transfected with the indicated plasmids: either pNKaf-GFP or pcDNA-GFP AND either pcDNA control vector or TSPyV circALTO plasmids. HSP90 serves as the loading control.
(E) IF images of 293T cells co-transfected with either pNKaf-GFP or pcDNA-GFP AND either pcDNA or TSPyV circALTO for 48 hours. Scale bar = 200 μm.
(F) Transcript level of GFP from 293T cells co-transfected with either pNKaf-GFP or pcDNA-GFP AND either pcDNA or TSPyV circALTO. *ACTB* served as the internal control. Error bars = SD from a single experiment. Results are representative of 2 independent experiments.

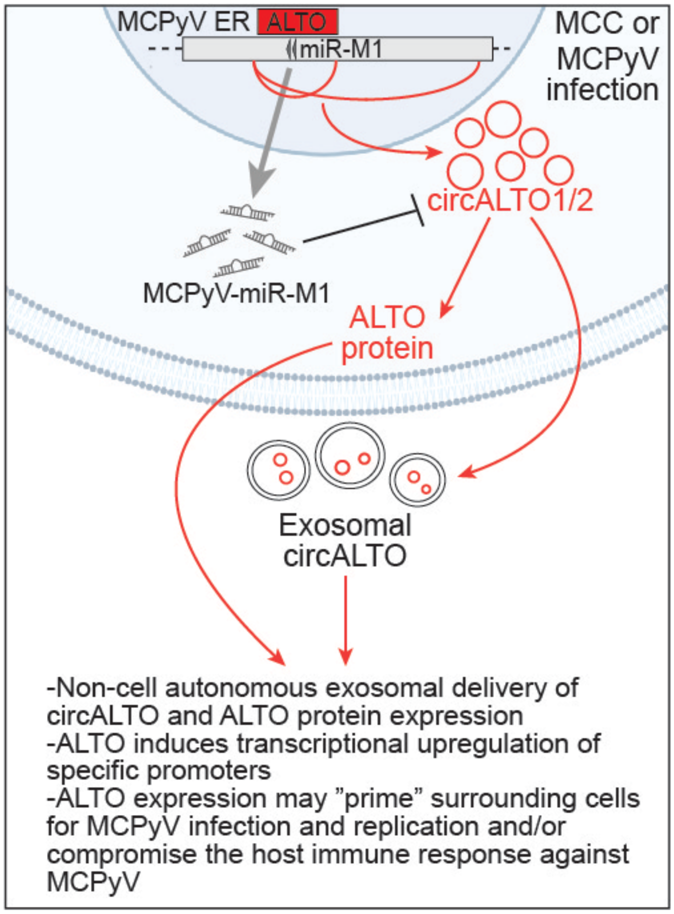


**Supplementary Figure S6. Graphical Abstract**

Model for the regulation and function of MCPyV circALTO.
